# Triphenylphosphonium is an effective targeting moiety for plants mitochondria

**DOI:** 10.1101/2025.02.20.638842

**Authors:** Shani Lazary, Gal Maman, Jenia Binenbaum, Mordechai Ronen, Iris Tal, Elon Yariv, Eilon Shani, Roy Weinstain

## Abstract

Small signaling molecule regulates key physiological processes in plants, often in a spatially distinct manner. However, current methods for applying small-molecules, endogenous or synthetic, in plants research lack spatial precision, limiting the ability to study and utilize their localized effects. Here, we validate triphenylphosphonium (TPP) as a mitochondrial targeting motif in plants. Using fluorescently labeled TPP conjugates in Arabidopsis thaliana, we demonstrate mitochondria-specific accumulation, even in the presence of plastids. This precise localization enables detailed imaging of mitochondria and mitochondrial DNA in living plants. We further exploit this targeting ability by developing a TPP-ciprofloxacin conjugate to selectively inhibit mitochondrial DNA gyrase, an enzyme involved in organellar DNA replication. Unlike free ciprofloxacin, which disrupts both mitochondrial and chloroplast DNA gyrase activity, the TPP-conjugate specifically targets mitochondrial gyrase, leading to slower plant growth without affecting chloroplast function. This targeted inhibition triggers a mitochondrial retrograde response, characterized by increased reactive oxygen species levels and the upregulation of stress-response genes in the nucleus. Our findings establish TPP as a reliable tool for mitochondrial targeting in plants and open avenues for both fundamental research and agricultural applications. By enabling organelle-specific manipulation in species not amenable to genetic engineering, TPP-based strategies have potential for advancing plant biology and precision agriculture.

## Introduction

Signaling small-molecules are instrumental in shaping plants physiology and adaptive growth.^1-4^ Most signaling molecules in plants regulate and influence multiple processes, having distinct molecular and phenotypic effects in different cells and even in cellular compartments (e.g. auxin^5-6^, gibberellins^7-8^, cytokinins^9-10^, hydrogen peroxide^11-12^, calcium^13-14^, nitrogen sources^15^, lipids^16-17^, and pathogenic molecules^18^). The diversity of functions each of them display, compounded by the differential responses they invoke in different locations, require techniques that operate at high resolution to dissect their distinct roles. Yet, in contrast to the remarkable progress made in manipulation of DNA, RNA and proteins, the ability to manipulate responses to small-molecules in plants has remained largely the same; they are invoked by indiscriminate application of the small-molecule to the whole plant, or at the organ level at most, which leads to undirected biological activity that is dependent solely on the molecule’s biodistribution pattern. By allowing small-molecules to freely distribute within the plant, the spatial context in which they often act is completely lost.

Although synthetic biology has proved useful for the biosynthesis of some endogenous, small organic molecules *in planta* with spatial and/or temporal control^19-20^, such genome manipulation is often accompanied by unintended developmental consequences and is an effort-intensive, *ad hoc* solution. Furthermore, the number of plant species that are amenable for genetic engineering is limited and the strategy is strictly restricted to those molecules for which a complete suite of biosynthetic enzymes is known and available. The same is true for many valuable synthetic small molecules in plant research, such as specific inhibitors and activity modulators, which are becoming increasingly more available thanks to progress in chemical-genomics screens.^21-22^ Such molecules can be applied in a conditional, dose-dependent and reversible manner, with the advantage of circumventing the limitations of lethality and functional redundancy inherent to mutants^23^, yet, their potential utility is far from being fully realized due to lack of ability to target them to specific locations. The long-standing inability to target bioactive molecules to specific sub-cellular locations also has implications in agriculture; only a fraction of an agrochemical applied in field reaches the site of action in crops^24-27^ (typically, a specific cellular compartment), resulting in inefficient utilization of resources and in significant environmental impact.

In recent years, several efforts have been made to facilitate targeted delivery of small-molecules in plants, either by utilizing light-mediated techniques^28-30^ or via chemical targeting motifs, to cells and organelles such as the chloroplast^31-32^ (Chl), vacuole^33^ and cell membranes^33^, highlighting the feasibility and the potential utility of this approach in both basic and applied plants research. We hypothesized that validating a generic mitochondria targeting motif in plants will engender the development of valuable tools to study this important organelle^34^ and its components in live, whole plants. Although several mitochondria targeting motifs have been characterized in mammalian cells, none has been comprehensively validated in plants. Out of these known mitochondria targeting motifs^35^, we chose to focus on triphenylphosphonium (TPP), mainly due to its broad scope of efficacy in organisms other than mammals^36-39^, its structural simplicity and its inherent hydrophobicity that could support efficient uptake into plants. The selective accumulation of TPP, and other similar cations, in mitochondria relies on its ability to diffuse through membranes and through the unique mitochondrial inner membrane potential (ΔΨm ∼ 120-180 mV), with the negative potential residing on the matrix side.^40^ However, unlike most other organisms, plants contain chloroplasts, a photosynthetic double-membrane organelle that exist in most plant cells in multiple copies. The envelope membrane potential in chloroplasts was measured to be -110 mV^41^, with the negative potential residing on the stroma side. The membrane potential of the thylakoids is reported to be 10-50 mV^42-43^, with the negative potential residing, again, on the stroma side. Thus, TPP accumulation in the chloroplast stroma might compete with its accumulation in the mitochondria. Indeed, several studies on isolated chloroplast have demonstrated the ability of TPP to penetrate into this organelle.^41, 44^

In this work, we evaluate the distribution pattern of TPP in whole, live plants via fluorescence labelling, confirming its selectivity to the mitochondria in this organism. We then show that TPP can be further leveraged to construct targetable tools to study this important organelle in plants.

## Results

### TPP selectively accumulates in plants mitochondria

The cellular distribution pattern of TPP was examined using fluorescently labeled conjugates, synthesized in three steps from TPP (Figure 1A, compounds **1**–**3**). The labeling fluorophores differ mostly in their electronic charge, intending to evaluate the effect of the cargo properties on parameters such as tissue penetration and distribution of the conjugate.

**Figure 1:**
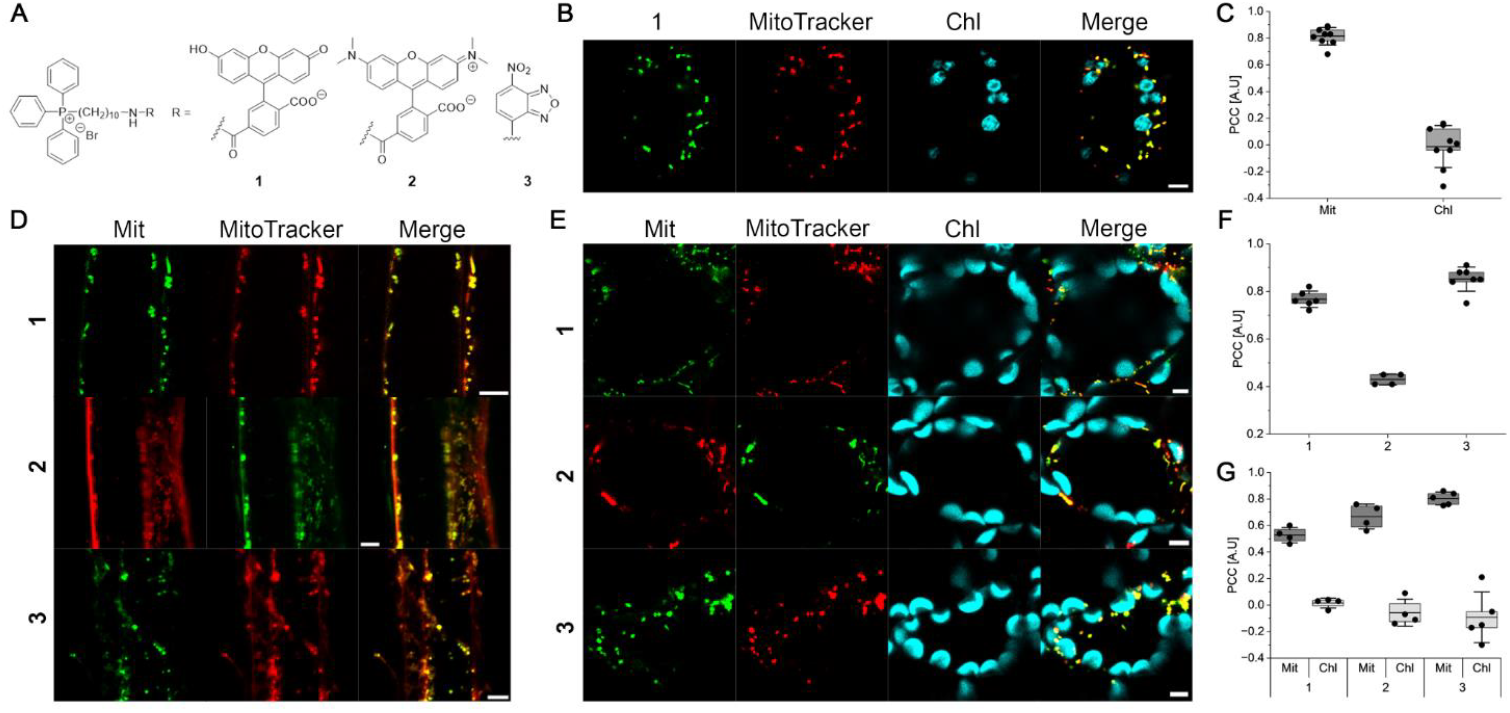
Fluorescent conjugates of TPP accumulate in mitochondria of *Arabidopsis thaliana* cell cultures and seedlings. **A)** Structures of synthesized TPP conjugates **B)** Representative confocal images of *Arabidopsis thaliana* cultured cells after incubation with **1** (25 μM, 30 min) and MitoTracker red (1 μM, 30 min). **C)** Box plot chart presenting co-localization between mitochondria (Mit) and chloroplasts (Chl) in cells using Pearson correlation coefficients (PCC) of non-thresholded images as shown in **B. D)** Representative confocal images of 5 days old *Arabidopsis thaliana* seedlings roots incubated for 3 h with MitoTracker red (1 μM) or green (10 μM) and the corresponding fluorescent probes **1** (25 μM) or **2** (10 μM) or **3** (10 μM). **E)** Representative confocal images of 5 days old *Arabidopsis thaliana* leaves. For **1**, seedlings were vacuum infiltrated for 10 min, then left for 3 h incubation with MitoTracker red. For **2** and **3**, seedlings were incubated for 3 h with MitoTracker red (1 μM) or green (10 μM) and the corresponding fluorescent probes **2** (10 μM) or **3** (10 μM). **F)** Box plot chart presenting co-localization analysis in roots for each probe and its co-responding MitoTracker using PCC of non-thresholded images as shown in **D. G)** Box plot chart presenting co-localization analysis in leaves for each probe and its co-responding MitoTracker and Chl natural fluorescence, using PCC of non-thresholded images as shown in **E**. For **B**,**D** and **E** bars represent 5 μm. For **C**,**F** and **G**, center lines represents the means and box limits indicate the 25^th^ and 75^th^ percentiles. Whiskers indicate ± SD.

In light-cultured *Arabidopsis thaliana* cells, conjugate **1** accumulated selectively in punctate structures that co-localized with the mitochondria marker MitoTracker red, and was absent from any other organelle, including from plastids (Figure 1B,C). Similar results were observed when wild-type (WT) *Arabidopsis thaliana* seedlings (Col-0) were incubated in solutions of **1**–**3**; the fluorescently labeled TPP derivatives accumulated exclusively in punctate structures that were co-labeled by either a green or a red MitoTracker marker, in both root and leaf cells (Figure 1D–G). In comparison, the respective free dyes showed mostly non-specific cellular distribution in these tissues (Figure S1). The identity of the punctate structures in which the fluorescently labeled TPP accumulated as mitochondria was further solidified by their co-localization with a GFP fusion of the mitochondria membrane protein UBIQUITIN SPECIFIC PROTEASE 27 (UBP27) in transgenic *Arabidopsis* seedlings (*p35S::UBP27-GFP-BLRP*, Figure 2A). Notably, the fluorescent signal of **2** was observed in the organelle interior (Figure 2B), suggesting it accumulates in the mitochondria matrix, in accord with the large negative membrane potential across the inner mitochondrial membrane.

**Figure 2:**
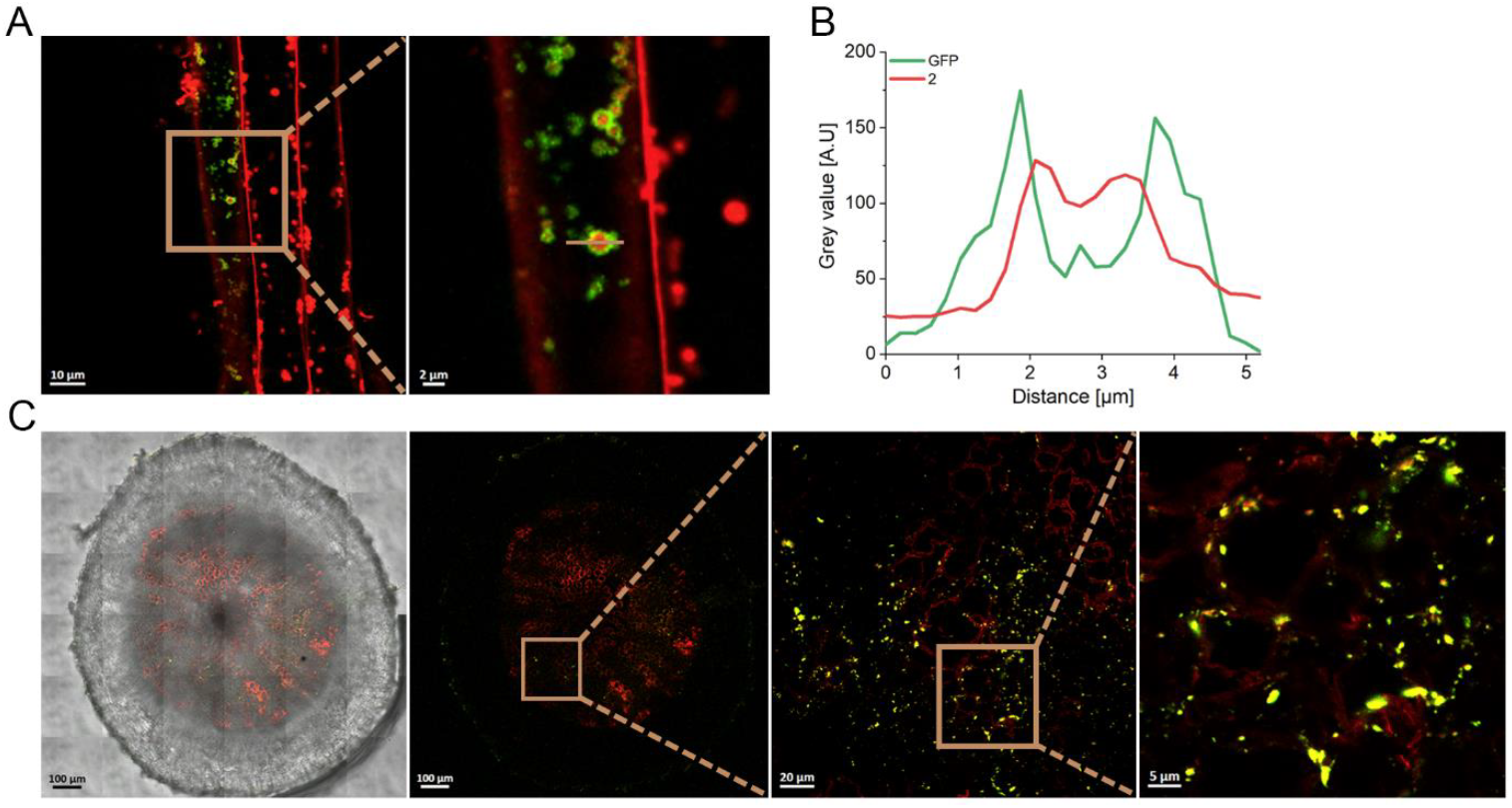
Fluorescent conjugates of TPP accumulate in *Arabidopsis thaliana* mitochondria matrix. **A)** Representative confocal images of 5 days old *p35S::UBP27-GFP-BLRP* transgenic *Arabidopsis thaliana* seedlings incubated with **2** (10 μM, 3 h). **B)** Line chart representing the grey value in the mitochondrion labeled in **A**, showing accumulation of **2** in a mitochondrion interior. **C)** Representative confocal images of a 1-month old *Arabidopsis thaliana* plants hypocotyl cross-section. Only the roots of the plants were treated for 16 h with **2** (10 μM) and MitoTracker green (10 μM), then hypocotyls were fixed in agarose, sectioned and imaged.

To assess the mobility of TPP conjugates *in planta* and their ability to reach deep tissues, only the roots of 1-month old *Arabidopsis* plants were dipped in a solution of **2** (10 μM) and MitoTracker green (10 μM) for 16 hours, followed by fixation and agarose embedment. Cross sections of the hypocotyl revealed that compound **2** translocated acropetally and accumulated in mitochondria of inner-layers cells (Figure 2C).

The effect of the cargo charge on TPP distribution pattern is worth noting; TPP conjugates with a negative-or a neutral-charge cargo (**1** and **3**, respectively) were highly specific to the mitochondria. However, ∼2.5-fold higher concentration of **1**, bearing a negatively charged cargo, was required to obtain a detectable signal *in planta* compared to the neutral-cargo bearing **3**, suggesting a lower uptake of the former. A cargo with a positive-charge (**2**) led to some association of the conjugate with the cell wall (Figure 1D, Figure 2A), presumably with the pectin-rich primary cell wall.^45^ Interaction of positively charged species with the cell wall is well documented.^46-48^ While these cellular structures (i.e. cell wall and mitochondria) can be easily differentiated visually, the effect should be considered when the cargo is non-fluorescent.

To evaluate the biodistribution of TPP conjugates in other plant species that are not readily amenable to genetic manipulations, compound **3** (10 μM) was applied to seedlings of *Lamium amplexicaule* (henbit), *Aegilops longissimi* (a wheat relative) and *Setaria viridis* (green foxtail) in solution and their hypocotyls (which contain chloroplasts) were consequently imaged. The results recapitulated those observed in *Arabidopsis*; selective accumulation of **3** in punctate structures that co-localized with MitoTracker red (Figure S2). *Aegilops longissimi* characteristics allowed for more comprehensive imaging, showing accumulation in different plant organs, such as root hairs and the hypocotyl (Figure S3).

Collectively, these data establish that TPP conjugates are readily taken up by plants, translocate, and accumulate in the mitochondria, including in photosynthetic organs, and suggest it can be utilized to localize molecules of interest selectively in the mitochondria of live plants. We therefore next evaluated the potential utility of TPP in plants for development of mitochondria-selective effectors.

### A TPP-Hoechst dye conjugate selectively labels mtDNA *in planta*

Mitochondrial DNA (mtDNA) plays important roles in diverse physiological processes and its direct imaging is an indispensable molecular tool for their studying.^49^ Several DNA-intercalating dyes are used for imaging of mtDNA by diffraction-limited microscopy, such as DAPI^50-51^, SYBR dyes^52-53^, and picoGreen^54-55^, including in plants.^52^ However, these dyes have a substantial drawback as they stain all DNAs of the cell, and therefore require an elaborate treatment or image analysis to avoid the significant fluorescence emanating from the abundant nuclear DNA (nuDNA). We speculated that conjugation of a fluorogenic DNA intercalator with a mitochondria-trappable entity might afford a “turn-on” fluorescent stain suitable for specific mtDNA imaging.

To test this concept, we synthesized a TPP conjugate that resembles the Hoechst 33342 dye (compound **4**, Figure 3A), one of the most prevalent DNA markers. The strong affinity of this dye family to DNA (binding constant of ca. 10^8^ M^-1^)^56^ has been previously leveraged for targeted delivery of conjugates to mammalian cells nuclei.^57-60^ Conjugate **4** exhibited a DNA-dependent fluorescence emission enhancement (Figure 3B), typical of the Hoechst dyes family^61^, validating that the conjugation to TPP does not interfere with the dye’s ability to intercalate within DNA. As expected, application of the free Hoechst dye 33342 to transgenic *Arabidopsis* seedlings expressing H2B-RFP (*35S::H2B-RFP*) as a fluorescent nuclear marker resulted in exclusive staining of nuDNA, as evident by the co-localization of the two fluorophores (Figure 3C). In comparison, following the application of **4** to similar seedlings, the fluorescence emission was observed exclusively in intracellular punctate structures and was completely devoid from the nucleus. The identity of these punctate structures as mitochondria was independently validated in transgenic seedlings expressing a genetically encoded mitochondria marker (*p35S::UBP27-GFP-BLRP*) (Figure 3D).

**Figure 3:**
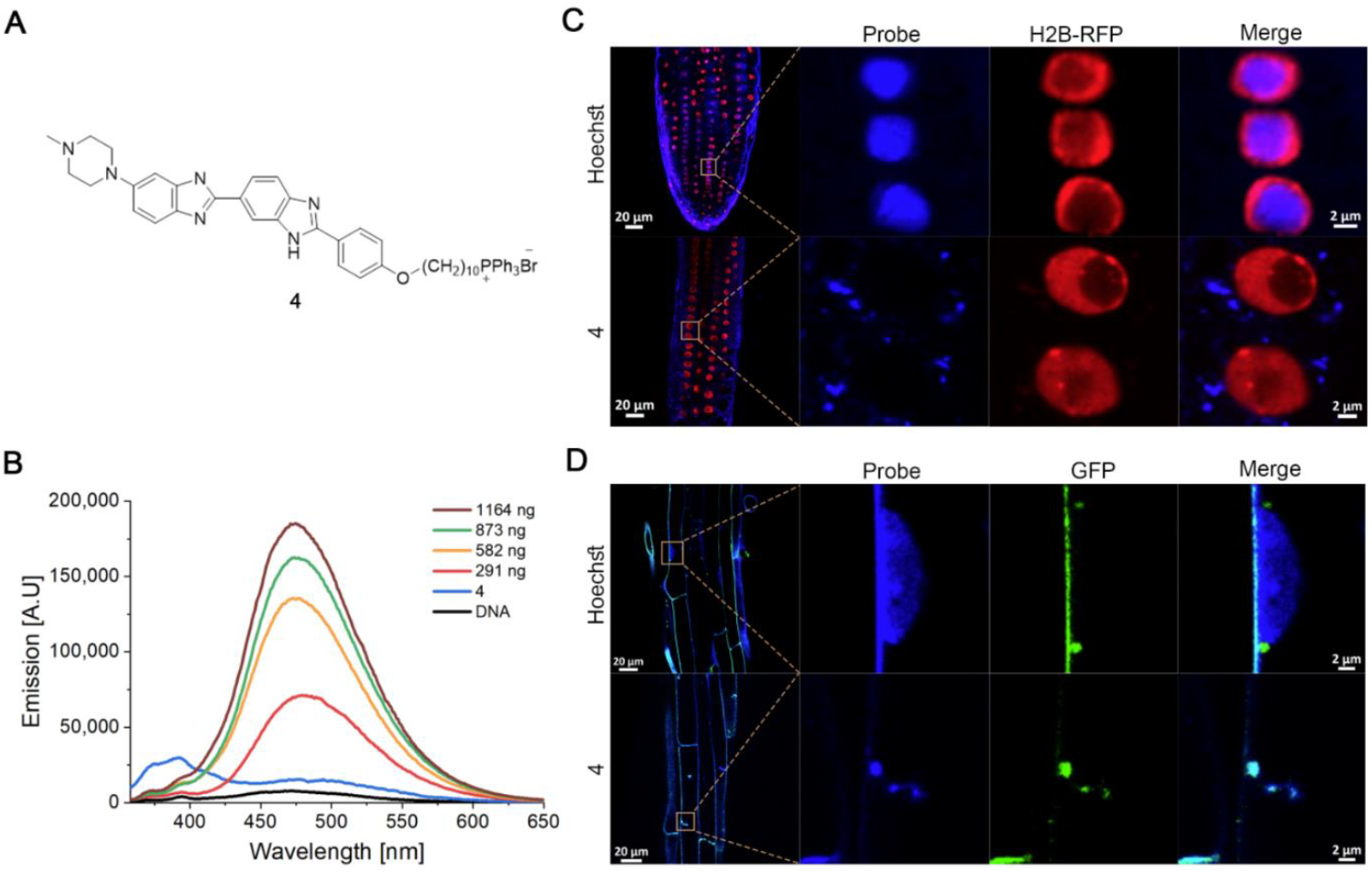
A TPP-Hoechst dye conjugate selectively stains mtDNA *in planta*. **A)** Structure of synthesized Hoechst dye 33342 conjugate with TPP **4. B)** Fluorescence emission of **4** with increasing amount of DNA following excitation at 348 nm. **C)** Representative confocal images of 5 days old *p35S:H2B-RFP* transgenic *Arabidopsis thaliana* seedlings incubated with Hoechst 33342 (2.5 μM, 3 h) or **4** (5 μM, 3 h). **D)** Representative confocal images of 5 days *p35S::UBP27-GFP-BLRP* transgenic *Arabidopsis thaliana* seedlings incubated with Hoechst 33342 (2.5 μM, 3 h) or **4** (5 μM, 3 h).

These results not only reinforce the specificity of TPP accumulation in plants mitochondria but also establish the ability of the conjugated cargo to access the mitochondria matrix and to further interact with endogenous entities within it.

### A TPP-ciprofloxacin conjugate selectively inhibits mitochondrial gyrase

To take further advantage of this capability, we sought to deliver an enzyme inhibitor specifically to the mitochondria matrix. Small molecule inhibitors are an indispensable tool for dissecting cellular functions. Nevertheless, they typically lack the spatial specificity that would allow them to differentiate between targets that are present in multiple cellular organelles. For example, DNA gyrase is a type-II topoisomerase that has key roles in DNA replication and transcription^62^ and that localizes in plants to both the chloroplast and the mitochondria.^63^ The specific functions of this enzyme in plants are not clearly understood as most eukaryotes do not express it. Previous studies have shown that knockout of the gyrase genes lead to embryo- or seedling-lethal phenotypes, impeding analysis of their function through traditional genetic tools.^63^ Quinolone and aminocoumarin antibiotics were shown to be effective inhibitors of plant gyrases^63-64^ and are even being explored as potential novel herbicides.^65^ However, both inhibitors lack the spatial specificity that would allow dissecting the differential roles of DNA gyrase in the chloroplasts and in the mitochondria. We hypothesized that a mitochondria-targeted ciprofloxacin (CFX), a known gyrase inhibitor of the quinolone family^66^, would distinguish between mitochondrial and chloroplast gyrases and help gain a better understanding of its mitochondria-specific roles.

To this end, we synthesized two derivatives of ciprofloxacin-TPP conjugates, differing in the functional group connecting the two units (**5** and **6**, Figure 4A). Both alkylation (**5**) and acylation (**6**) of the piperazine secondary amine were previously utilized for derivatization of CFX and were shown to not significantly interfere with its mechanism of inhibition.^67^ Computational docking experiments showed no significant differences in the estimated binding affinities and that the binding modes were conserved for the CFX part (Figure 4B–D). The aliphatic addition lies near the DNA coil and does not cause allosteric clashes. These results were observed for both the bacterial gyrase (*Staphylococcus aureus*, 2XCS crystal structure) and for the two *Arabidopsis* gyrase complexes (*At*GyrA - At3g10690 and chloroplast *At*GyrB - At3g10270 or mitochondrial *At*GyrB - At5g04130, using alpha-fold2 structural predictions based on the 2XCS experimental data), which show high similarity in their sequence to the bacterial enzyme and that were shown to bind CFX.^64^ Both compounds showed ∼30% retention of gyrase inhibition activity compared to free CFX against the *E. coli* gyrase in a relaxation of supercoiled DNA assay *in vitro*, with compound **6** showing higher potency at higher concentrations (Figure 4E). Therefore, we focused further efforts on the more potent compound **6**.

**Figure 4:**
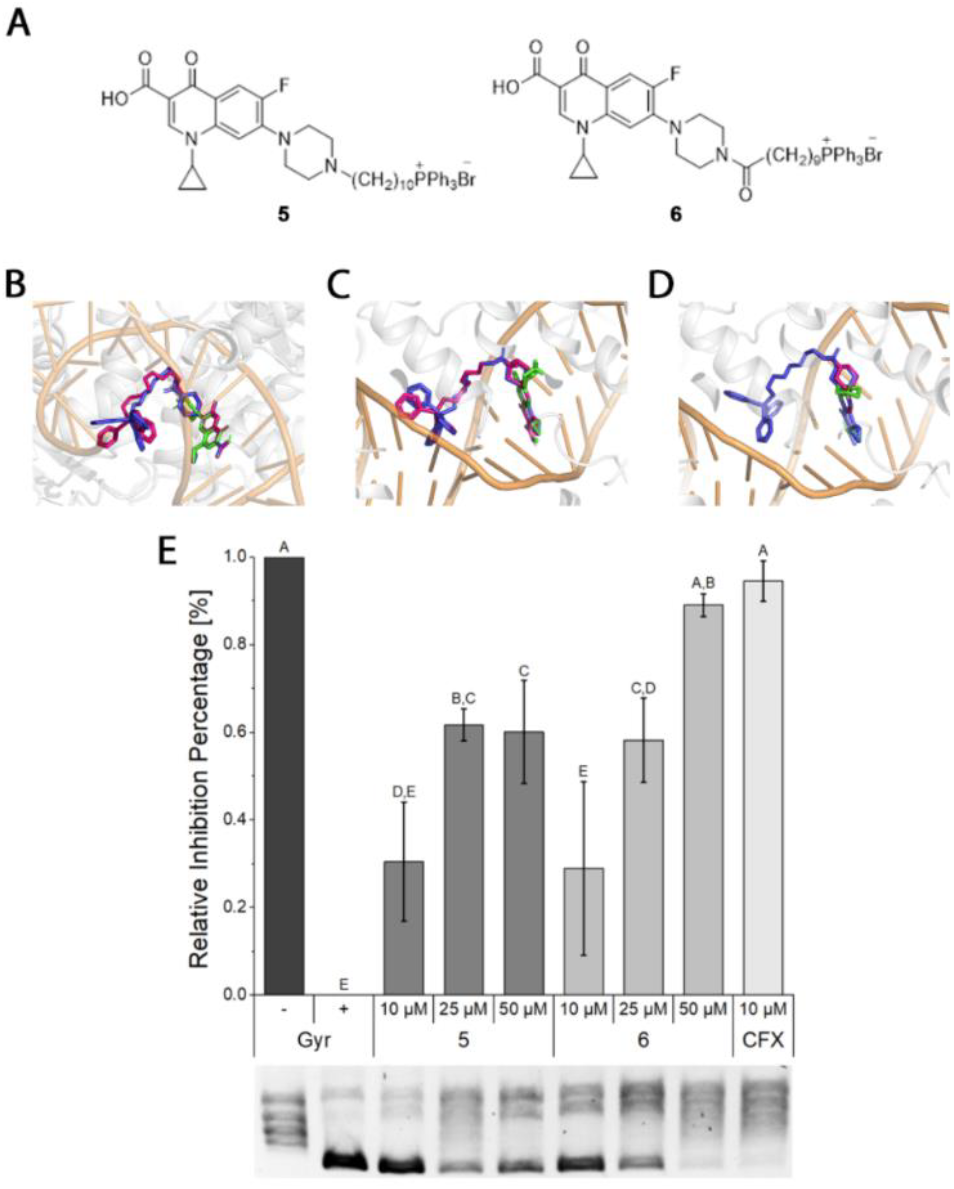
TPP conjugates of CFX retain inhibitory activity against DNA gyrase. **A)** Structures of synthesized CFX-TPP conjugates. **B**–**D)** Three dimensional representations of CFX (green), CFX-TPP conjugates **5** (magenta) and **6** (blue) molecular docking poses in: **B)** *S. aureus*, gyrase crystal structure (PDB ID: 2XCS) or alpha-fold2 models of *Arabidopsis thaliana* **C)** chloroplasts gyrase and **D)** mitochondrial gyrase **E)** Gyrase inhibition assay and bar chart of gyrase relative inhibition. Whiskers indicate ± SD. Significance was determined by one-way ANOVA analysis with Tukey’s honestly significant difference (HSD) post-hoc test. Different capital letters (A–E) indicates a significant difference at a p.v < 0.05.

A root elongation assay performed on 5-days old WT Arabidopsis (Col-0) seedlings showed that **6** has an EC_50_ value ∼10-times higher than CFX (16.2 versus 1.4 μM, respectively, Figure 5A and SI Figure 4A,B), in agreement with the results obtained for inhibition of the *E. coli* gyrase. To assesses the specificity of **6** towards the mitochondrial gyrase *in planta*, 14-days old *Arabidopsis* plants were sprayed with free CFX or **6** (10 or 50 μM, respectively) and subsequently monitored for 14 days. Within the period of observation, plants treated with CFX maintained a similar growth rate to that of untreated ones (Figure 5B), but showed an abnormal leaves shape and progressive decolouration (SI Figure 4C–E), indicative of chloroplasts dysfunction. Correspondingly, measured photosynthetic parameters, such as photosystem II quantum yield, showed a significant drop in comparison to non-treated plants (Figure 5C and SI Figure 4F–H). In contrast, plants treated with **6** maintained photosynthetic parameters similar to those of untreated plants but were significantly smaller, presumably indicative of mitochondria dysfunction below a minimal threshold required to support growth.^68^ The fact that the phenotypic manifestation of free CFX effect is observed mostly via chloroplast functions could be a result of its distribution being skewed towards this organelle or that chloroplasts are more sensitive to its effect than the mitochondria. In agreement with the above results, transmission electron microscopy (TEM) images of plants treated with CFX revealed structurally deformed chloroplasts with altered thylakoid arrangements, as previously reported^64^, and disrupted mitochondria morphology, while **6** treated plants showed normally looking chloroplasts and structurally deformed mitochondria (Figure 5D).

**Figure 5:**
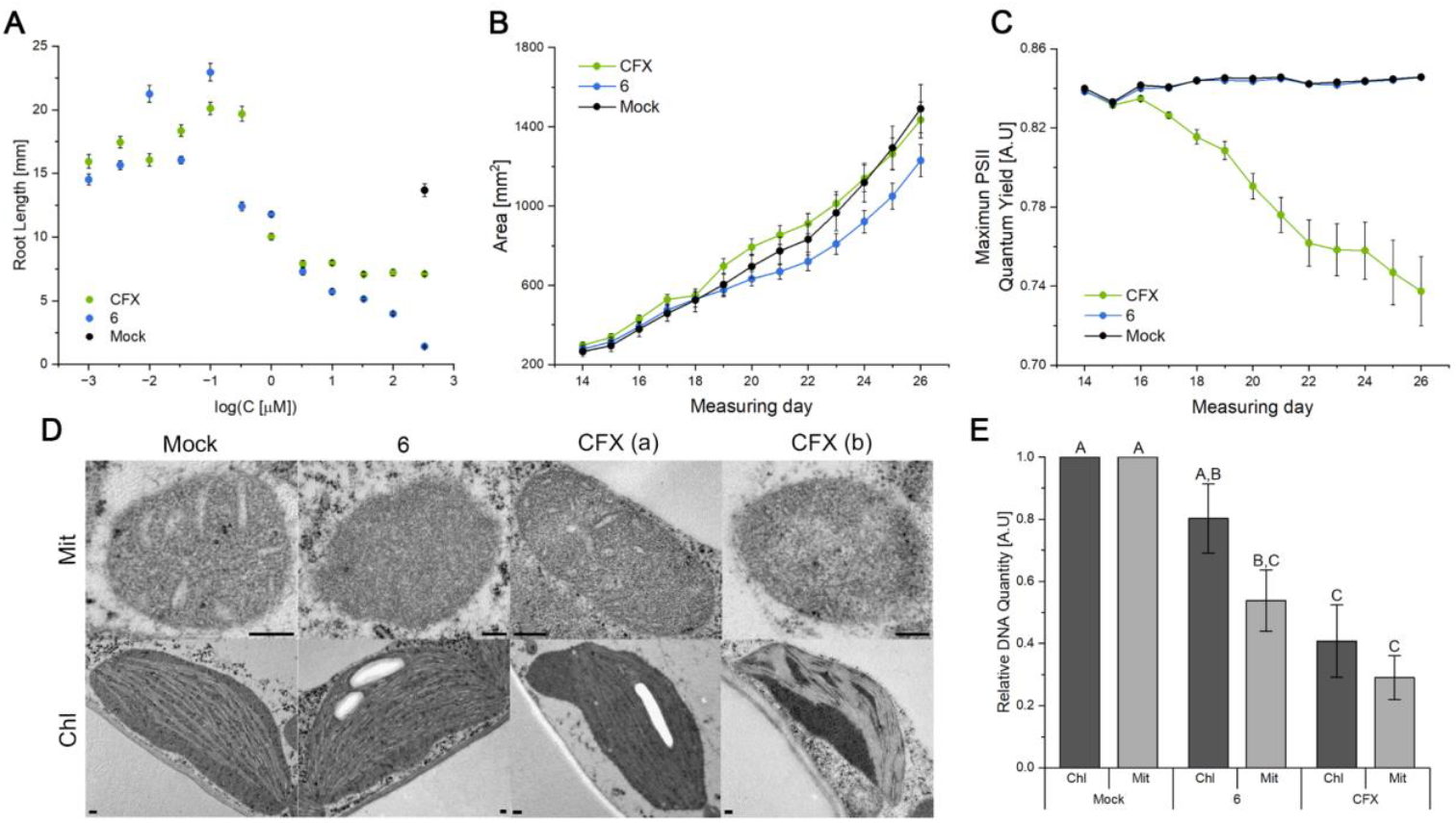
A CFX-TPP conjugate inhibits *Arabidopsis thaliana* mitochondrial DNA gyrase *in planta*. **A)** Root elongation essay of 5-days old *Arabidopsis thaliana* seedlings grown for 5 days on indicated concentrations of CFX or **6**. Each data point represents an average of n ≥ 8 plants. **B**–**C)** Measurement over 2 weeks of *Arabidopsis thaliana* plants rosette leaves area (**B**) and maximal PSII quantum yield (**C**), after treated twice (on days 10 and 12) with CFX (10 µM) or **6** (50 µM) by spray. Each data point represents the mean of 20 plants. **D)** TEM images of organelles from 2 weeks old *Arabidopsis thaliana* plants treated with CFX (10 µM) or **6** (50 µM) for 24h. Bars represent 200 nm. **E)** Relative DNA expression, quantified by qRT-PCR, of 5 days old *Arabidopsis thaliana* seedlings treated for 24 h with CFX (10 µM) or **6** (50 µM). Significant was determined by one-way ANOVA analysis with Tukey’s honestly significant difference (HSD) post-hoc test. Different capital letters (A–C) indicate a significant difference at a p.v < 0.05. For **A, B, C**, and **E** whiskers indicate ± SE.

A previous study has found that following treatment with CFX, there is a reduction in the number of both chloroplasts and mitochondria in wild-type plants.^64^ To evaluate the effect of the targeted CFX, 5-days old *Arabidopsis* plants were treated with either CFX or **6** (10 or 50 μM, respectively) and the amount of mitochondrial and chloroplast DNA (as a proxy for organelles amounts) was quantified 24 hours later with respect to the amount of nuclear DNA using qPCR for two genes in each organelle (Figure 5E). Treatment with CFX led to significant reduction in the amount of both mitochondrial and chloroplast DNA while treatment with **6** led to significant reduction in mitochondrial DNA and non-significant (∼20%) reduction in the amount of chloroplast DNA. These results can suggest that some mitochondria-targeted CFX ends up in the chloroplast, directly affecting this organelle, or, alternatively, that the reduction observed in the amount of chloroplasts is a downstream effect of reduction in the amount of functional mitochondria.^69^ Collectively, these results suggest that compound **6** effectively inhibits DNA gyrase specifically in plants mitochondria.

Many stress conditions in plants lead to elevated reactive oxygen species (ROS) levels, which in turn, activate various stress-response pathways.^4^ We hypothesized that interfering with mitochondrial replication via gyrase inhibition should lead to some form of a stress response. Indeed, when **6** was injected to *Nicotina benthamiana* leaves, a significant increase in ROS level, as measured by 2’,7’-dichlorodihydrofluorescein diacetate (DCF) fluorescence intensity, was observed 3 hours post-treatment in the mitochondria but not in chloroplasts, compared to leaves injected only with water (Figure 6A,B). This suggests that a mitochondrial mechanism^4^ senses the defunct activity of the gyrase, or its downstream effects, and responds by increasing ROS levels. Moreover, a significant increase in the transcription of ARABIDOPSIS NAC DOMAIN PROTEIN 13 (ANAC013) transcription factor and the downstream ALTERNATIVE OXIDASE 1A (AOX1a) gene was observed 3 hours post-treatment with **6** (Figure 6C). Both genes are common markers of mitochondrial stress responses that are mediated by ROS signaling.^70^

**Figure 6:**
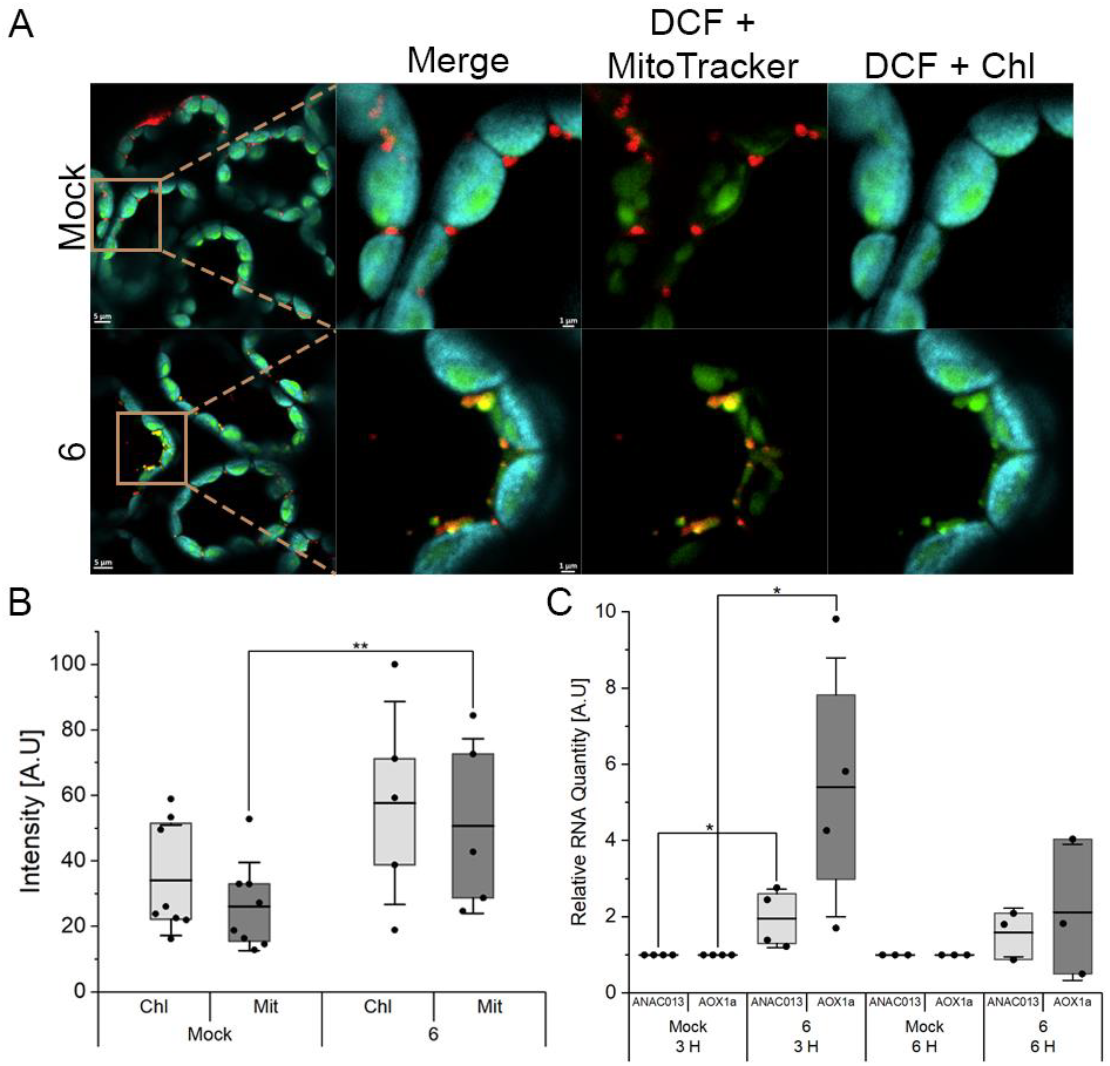
Inhibition of mitochondria DNA gyrase leads to nuclear stress response. **A)** Representative confocal images of 2 weeks old *Nicotiana benthamiana* leaves treated with **6** (50 μM) or DDW and incubated with DCF (50 µM) and MitoTracker (1 µM). **B)** Box plot presenting the intensity of DCF. Each point represents mean fluorescence intensity of 5 organelles randomly selected. **C)** Box plot presenting the relative RNA expression levels of ANAC013 and AOX1a transcripts, quantified by qRT-PCR, of 5 days old Arabidopsis seedlings treated for 3 h or 6 h with **6** (50 µM). For **B** and **C**, center lines represent the means and box limits indicate the 25^th^ and 75^th^ percentiles. Whiskers indicate ± SD. Significant was determined by one-tailed Welch *t*-test, * indicates a significant difference at a p.v < 0.05, ** indicates a significant difference at a p.v < 0.01.

## Discussion

The key outcome of this study is the validation of TPP as a reliable mitochondrial targeting motif in plants. This finding is particularly significant given the unique challenges posed by plant cells, which contain multiple organelles with overlapping structural properties, such as mitochondria and chloroplasts, and various barriers, such as the cell wall, that are not present in mammals. The observed selectivity suggests that TPP-based strategies could overcome the limitations of traditional small-molecule applications, which often lack spatial specificity and lead to off-target effects. Other mitochondria-targeting motifs are known^35^, but await further validation in plants.

The study highlights the potential of TPP conjugates for the development of mitochondria-selective probes and effectors. The successful conjugation of TPP to a Hoechst dye, resulting in specific labeling of mtDNA *in planta*, should provide a mean to study mitochondrial processes without signal interference from nuclear DNA, which is a common challenge when using traditional DNA-binding dyes. The ability to visualize mtDNA in live, whole plants with minimal background interference will enable more detailed investigations into its roles in plant development and stress responses.^34^ Moreover, to the best of our knowledge, this is the first specific mtDNA-visualizing agent reported in any organism and is therefore a valuable addition to the growing toolkit of mitochondria imaging probes.^71^

The TPP-CFX conjugate, which selectively inhibited mitochondrial DNA gyrase, represents an advancement in the ability to dissect organelle-specific functions. A similar approach could be applied to study other enzymes that reside in the mitochondria matrix but also in other organelles, such as superoxide dismutase^72^, aconitase^73^ and glutamine synthase.^74^ The differential effects observed between free ciprofloxacin and the TPP-conjugated version underscore the importance of spatial specificity in small-molecule applications. By restricting the activity of ciprofloxacin to mitochondria, we unveiled phenotypic and molecular effects that arise specifically from inhibiting DNA gyrase in this organelle, mostly avoiding confounding effects from gyrase chloroplast malfunction. The slower growth phenotype observed in seedlings treated with mitochondria-targeted ciprofloxacin is in accord with mitochondria dysfunction.^75-77^ Reproductive tissues are typically the ones most severely impaired by mitochondrial dysfunction, but vegetative development is also often affected, manifesting in slow growth and overall stunted stature.^78^ Mutants in other DNA replication machinery components, such as the mitochondrial polymerase IB, show a similar phenotype.^79^ Nevertheless, knockout of the mitochondrial *GyrB2* unit results in much more deleterious effects.^64^ This might suggest that mitochondrial gyrase activity is more vital in the first phases of germination and seedling development then in later ones, or simply reinforce the importance of a properly functioning mitochondria in these stages.^78^ Inhibition of the mitochondrial DNA gyrase was found to elevate ROS levels in the this organelle and to upregulate the transcription of ANAC013, a known mediator of mitochondria retrograde regulation (MRR)^80^, as well as AOX1a, by far the most commonly used indicator of mitochondrial retrograde response.^81-82^ This suggests that improper function of the mitochondrial DNA gyrase is signaled to the nucleus through the endoplasmic reticulum, presumably via ROS as an MRR-triggering signaling molecule.^83-85^

In conclusion, this study demonstrates a novel approach for controlling the bioactivity of small-molecules in plants with high spatial resolution by utilizing a targeting chemical moiety. It establishes TPP as a versatile mitochondrial targeting motif in plants, with potential for developing organelle-specific probes and effectors. The successful application of TPP-based conjugates in various plant species, especially ones that are not amenable to genetic engineering, suggests that this approach could be broadly applicable, offering new avenues for both basic research and agricultural innovation. Finally, the potential for cross-application of this approach to other organelles^86^ should allow for an expanded toolkit to dissect and manipulate the functions of distinct organelles.

## Data availability

All data supporting the findings of this study are available within the paper and its Supplementary Information.

## Author contributions

R.W. and S.L. conceived the project and R.W. supervised the project. S.L. designed and performed experiments and analyzed the data. G.M. and M.R. performed qPCR experiments. G.B performed cross sections samples and I.T. constructed the *p35S::UBP27-GFP-BLRP* transgenic lines under the supervision of E.S. M.R. designed and performed phenotypic experiments. E.Y performed computational predictions and docking experiments.

## Acknowledgements

We would like to thank E. Shani for providing *Arabidopsis thaliana* (Columbia ecotype) cell culture and *Arabidopsis thaliana p35S::H2B-RFP*^2^ transgenic line; A. Avni for providing *Nicotiana benthamiana* plants and maintaining the *Arabidopsis thaliana* cell culture; N. Ohad for providing *Lamium amplexicaule* seeds and N. Sade for providing *Aegilops longissimi* and *Setaria viridis* seeds. We thank L. Pizarro for her confocal training. We also want to thank A. Kessel for his help preparing *Staphylococcus aureus* (2XCS) and *Arabidopsis thaliana GyrA* and *GyrB* DNA sequences for computational modeling and H. Failayev for her help in optimizing the models’ visuals. Lastly, we thank V. Holdengreber at the Rosalie and Harold Rae Brown Cancer Research Core Facility for the TEM microscopy operation.

