## SUPPLEMENTARY INFORMATION for "Triphenylphosphonium is an effective targeting moiety for plants mitochondria"

**Synthesis of 6** 21
**Synthesis of 7** 21
**$^{1}$H- and $^{13}$C-spectra NMR** 23
**References** 34

### Supporting Figures

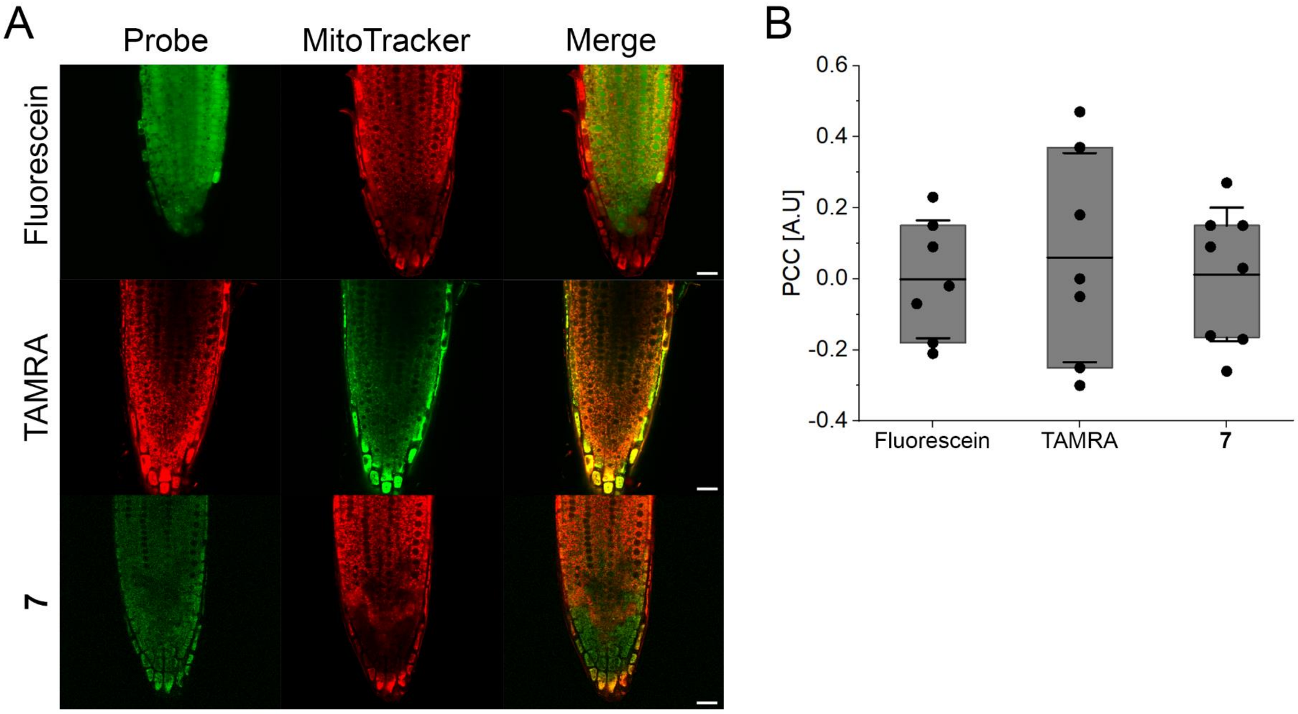

**Figure S1: Non-targeted fluorophores do not accumulate in mitochondria. A)** Representative confocal images of 5 days old *Arabidopsis thaliana* seedlings roots incubated for 3 h with MitoTracker red (1 µM) or green (10 µM) and the corresponding free fluorophores; Fluorescein (25 µM) or TAMRA (10 µM) or **7** (10 µM). Bars represent 20 µm. **B)** Box plot chart presenting co-localization analysis in roots for each fluorophore and its compatible MitoTracker using PCC of non-thresholded images as shown in **A**. Center lines represent the means and box limits indicate the 25$^{th}$ and 75$^{th}$ percentiles. Whiskers indicate ± SD.

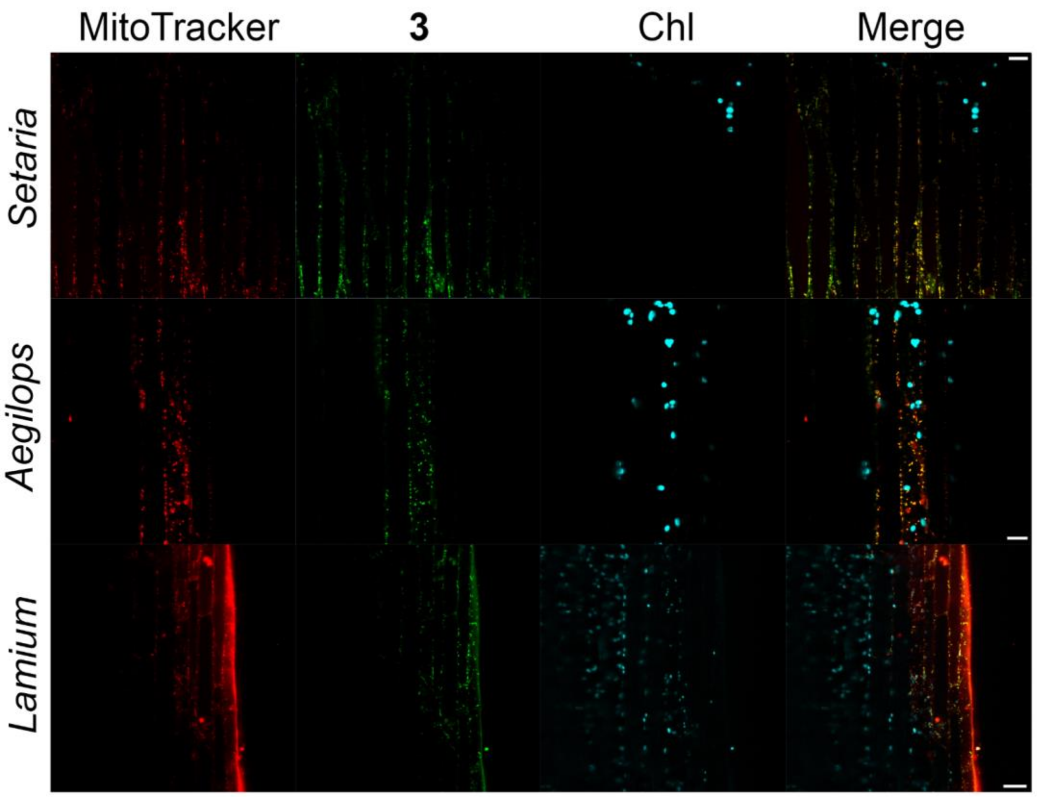

**Figure S2: Fluorescently labeled TPP localize to mitochondria in various plant species.** Representative confocal images of 5-7 days old *Setaria viridis*, *Aegilops longissimi and Lamium amplexicaule* seedlings roots incubated for 3 h with MitoTracker red (1 µM) and **3** (10 µM). Bars represent 10 µm.

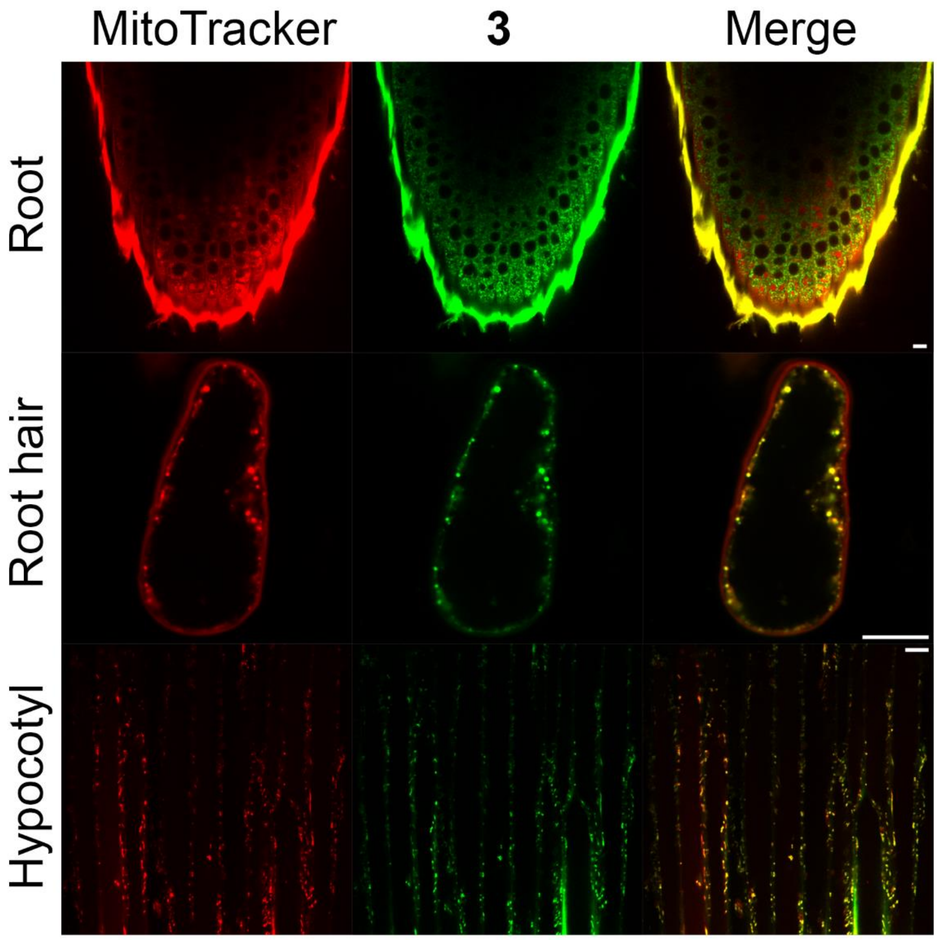

**Figure S3: Fluorescently labeled TPP localize to mitochondria in multiple organs of *Aegilops longissimi*.** Representative confocal images showing different organs of 5-7 days old *Aegilops longissimi* seedlings roots incubated for 3 h with MitoTracker red (1 µM) and **3** (10 µM). Bars represent 10 µm.

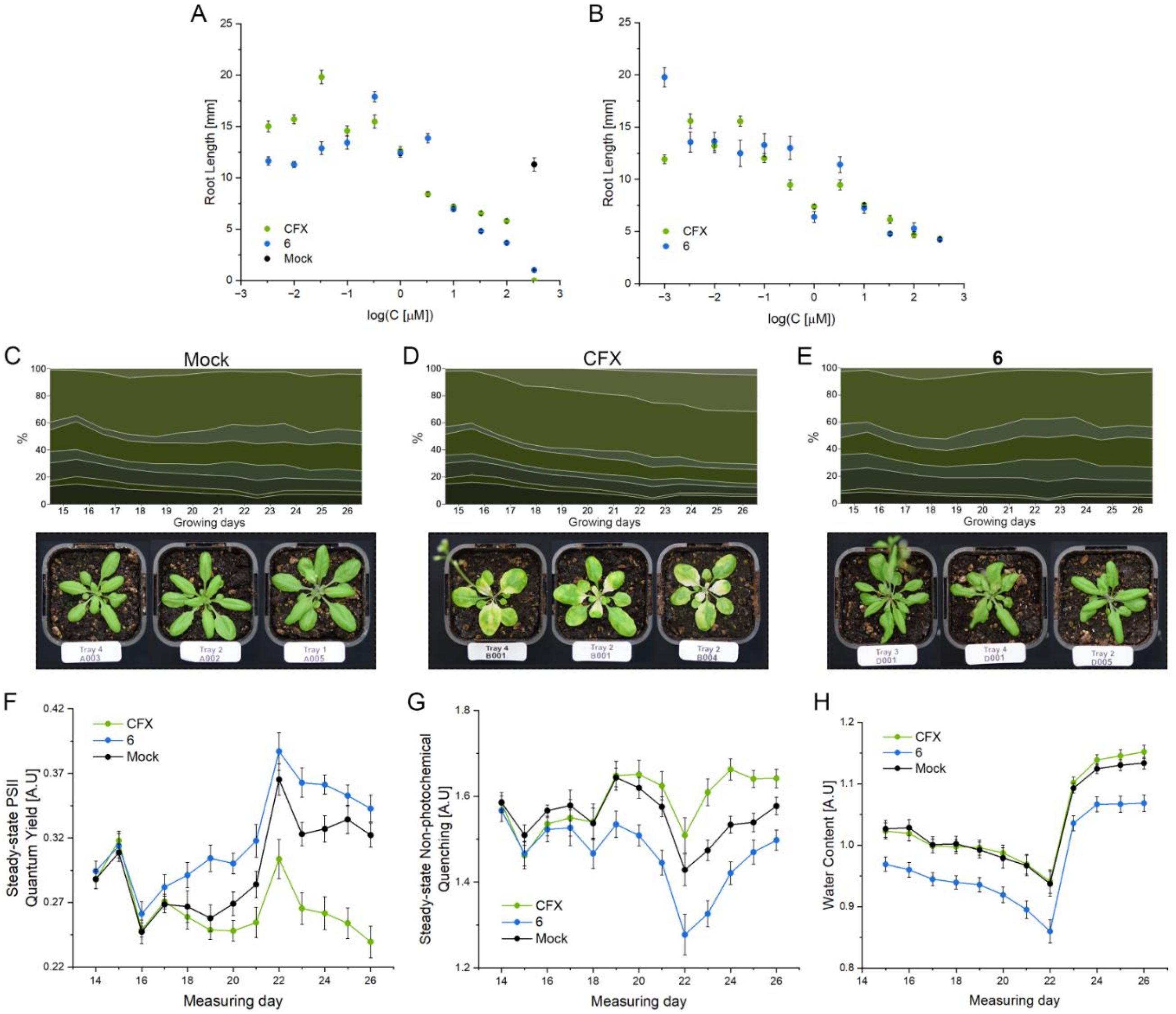

**Figure S4: A CFX-TPP conjugate inhibits *Arabidopsis* mitochondrial DNA gyrase *in planta*. A-B)** Two additional repeats for the root elongation essay. 5-days old *Arabidopsis thaliana* seedlings were grown for 5 days on indicated concentrations of CFX or **6**. Each data point represents an average of n ≥ 8 plants. **C-E)** Measurements of *Arabidopsis thaliana* plants color composition from 8 pigments over 2 weeks, after treated twice (on days 10 and 12) with mock (**C**) CFX (**D**, 10 μM) or **6** (**E**, 50 μM) by spray, alongside representative plants. Images were taken on the last measuring day. **F-G)** Measurement over 2 weeks of *Arabidopsis thaliana* plants steady-state PSII quantum yield (**F**), steady-state non-photochemical quenching (**G**) and water content (**H**), after treated twice (on days 10 and 12) with CFX (10 μM) or **6** (50 μM). Each data point represents the mean of 20 plants. For **A**, **B**, and **F-H**, whiskers indicate ± SE.

### Biological Methods

#### Plant material and growth conditions

*Arabidopsis thaliana* (Columbia ecotype) cell culture[1] was kindly provided by Prof. Eilon Shani's Lab and was kindly maintained in Prof. Adi Avni's Lab.

All *Arabidopsis thaliana* lines used in this work are Colombia background (Col-0 ecotype, Salk Institute La Jolla, CA, USA). Sterilized seeds were plated on Murashige & Skoog (MS) x 0.5 (Duchefa Biochemic) medium containing 1% w/v sucrose and 0.5% w/v plant agar (Duchefa Biochemic), pH was adjusted to 5.6-5.8 with 1 M KOH. Plates with seeds were stratified for 48 hours at 4 °C then transferred to growth chamber (Percival CU41L5) at 21 ˚C, 100-120 $\mu Em^{-2}S^{-1}$ light intensity under long-day conditions (16 h light/8 h dark).

For leaf imaging, plants phenotypic characterization and TEM experiments *Arabidopsis thaliana* seeds were sown on wet soil, stratified for 48 hours at 4 °C then transferred to growth room at 21 °C under long-day conditions.

*Arabidopsis thaliana p35S::H2B-RFP*[2] transgenic seeds were kindly provided by Prof. Eilon Shani's Lab.

*Nicotiana benthamiana* plants were kindly provided and maintained by Prof. Adi Avni's Lab. Seeds were sown on wet soil and transferred to growth room at 25 °C under long-day conditions.

*Lamium amplexicaule* seeds were kindly provided by Prof. Nir Ohad's Lab. *Aegilops longissimi* and *Setaria viridis* seeds were kindly provided by Dr. Nir Sade's Lab. *Lamium amplexicaule*, *Aegilops longissimi* and *Setaria viridis* seeds were sown on wet Watman Paper placed in plastic magenta box that left in growth room at 25 °C under long-day conditions.

#### Cloning and expression of GFP in *Arabidopsis thaliana's* mitochondria

*Arabidopsis thaliana p35S:UBP27$^{1-71}$-GFP-BLRP* were constructed as follow: *p35S:UBP27$^{1-71}$-GFP-BLRP* constructed utilizing Gibson method (NEB-E2611). The first 71 amino acids of UBP27 (AT4G39370)[3] were amplified from cDNA:

F primer: tctagtcgacctgcaggcggccgcaATGGTTTCTAGAAGAGGCTC.

R primer: ctcacagcggccgcAAAAGAATCGTCTCCGGAG.

eGFP- BLRP amplified from NTF[4]:

F primer: agacgattcttttgcggccgctGTGAGCAAGGGCGAGGAG.

R primer: ggagaccggcacactggccatatcgTCAAGATCCACCAGTATCCTCATG.

Fragments were assembled into pK2GW7 backbone and linearized with SpeI & BstXI.

**Seed sterilization**

*Arabidopsis thaliana* seeds from all lines were sterilized by vapor-phase sterilization (chlorine fumes). Seeds were placed in Eppendorf tubes inside a sealed desiccator, in the presence of 4% 32% hydrochloric acid in 11% sodium hypochlorite. After 2-3 h, seeds were taken out and left to ventilate for 30 min in biological hood.

*Aegilops longissimi* seeds were sterilized by incubation in 3% sodium hypochlorite solution for 3 min. the sterilized seeds were then washed 5 times with double distilled water (DDW).

*Lamium amplexicaule* and *Setaria viridis* seeds were sterilized by incubation in 3% sodium hypochlorite solution for 1 min. The sterilized seeds were then washed 5 times with DDW.

**General imaging**

All samples were imaged on a laser scanning confocal microscope (Zeiss LSM 780 inverted microscope), with 63X/1.15 W or 40X/1.2 W objective lenses with TPMT module for bright field. Hoechst 33342 and **4** were excited with 405 nm laser; emissions were collected between 410 nm to 556 nm. **1**, **3**, MitoTracker green and *p35s::UBP27-GFP-BLRP|BirA* transgenic lines were excited with a 488 nm laser; emissions were collected between 493 to 556 nm. DCF was excited with 488 nm laser; emission was collected between 501 nm to 576 nm. **2** and MitoTracker red CMXRos were excited with 561 nm laser; emissions were collected between 570 nm to 632 nm. *p35s:H2B-RFP* transgenic plants were excited with 561 nm laser; emission was collected between 582 nm to 754 nm. Chlorophyll was excited with 633 nm laser; emission was collected between 647 nm to 721 nm.

**Fluorescent probes imaging**

*Arabidopsis thaliana*

Cell culture was incubated with MitoTracker red (1 μM) and **1** (25 μM) for 30 min, then washed twice with DDW, mounted on a slide and imaged.

Roots: Seedlings were grown on agar as described in "Plant material and growth conditions" section. At 5-7 days seedlings were incubated in liquid MS with the inspected probes: MitoTracker red (1 μM) and **1** (25 μM); MitoTracker red (1 μM) and **3** (10 μM); MitoTracker green (10 μM) and **2** (10 μM); MitoTracker red (1 μM) and fluorescein (25 μM); MitoTracker red (1 μM) and **7** (10 μM); MitoTracker green (10 μM) and TAMRA (10 μM). The seedlings were left for 3 h, then washed twice with DDW and taken for imaging.

For all probes, co-localization analysis was done using Coloc2 function in Fiji software. Non-corrected and non-thresholded images of probes and respective MitoTracker were loaded to the software.

Pearson correlation coefficients (PCC) and Manders correlation coefficients above the autothreshold (tM1 and tM2) were determined from ROI ($n \geq 5$).

*p35S::UBP27-GFP-BLRP* transgenic *Arabidopsis thaliana* seedlings were grown on agar as described in "Plant material and growth conditions" section. At the 5-7 days, seedlings were incubated for 3 h in liquid MS with **2** (10 μM), then washed twice with DDW and taken for imaging. A line was drawn over an individual mitochondrion and grey values were measured using FIJI.

Leaves**:** **1** - Seedlings were placed in 3 mL wells containing **1** (25 μM) in liquid MS and were vacuum infiltrated for 10 min, then MitoTracker red (1 μM) was added and the seedlings left for incubation. After 3 h seedling were washed twice with DDW and taken for imaging.

**2** and **3** - *S*eedlings were grown in plates as described in "Plant material and growth conditions" section. At 5-7 days seedlings were incubated in liquid MS with the inspected probes: MitoTracker red (1 μM) and **3** (10 μM) or MitoTracker green (10 μM) and **2** (10 μM). The seedlings were left for 3 h, then washed twice with DDW and taken for imaging.

*Lamium amplexicaule*, *Aegilops longissimi* and *Setaria viridis*

Seedlings were grown in magenta boxes as described in "Plant material and growth conditions" section. At 5-7 days, seedlings were incubated for 3 h in liquid MS with MitoTracker red (1 μM) and **3** (10 μM), then washed twice with DDW and taken for imaging.

**Cross sections**

*Arabidopsis thaliana* cross sections were done as described previously by Skopelitis *at el.*[5] Briefly, a 3-weeks old Arabidopsis plant was incubated for 12 h in 0.5% liquid MS containing MitoTracker green (10 μM) and **2** (10 μM). The plant hypocotyl was fixed in 4% paraformaldehyde (PFA) dissolved in 1x PBS supplemented for 60 min. Fixed tissues were washed twice (10 min per wash) in 1× PBS and were then transferred to ClearSee solution. After 12 h the tissues were embedded in 8% Low Melting Agarose (Invitrogen) for 15 min and were then sectioned using a VT1000S vibratome (Leica) to generate 100 μm sections. Sections were then taken for imaging.

**Emission of 4 in the absence or presence of DNA**

Total DNA (extracted from *Arabidopsis thaliana*, as described in the "Relative DNA expression" section) was added incrementally to a cuvette containing **4** (5 μM) in DDW. Fluorescence emission spectrum ($\lambda_{ex}$ = 348 nm, $\lambda_{em}$ = 358-650 nm) was recorded after each addition. Data was processed using OriginPro software.

**Imaging of 4 in *Arabidopsis thaliana*'s roots**

*p35S:H2B-RFP* or *p35S::UBP27-GFP-BLRP* transgenic *Arabidopsis thaliana* seedlings were grown as described before. At 5-7 day, seedlings were incubated for 3 h in liquid MS with Hoechst 33342 (2.5 μM) or **4** (5 μM), washed twice with DDW, and subsequently imaged.

**Modelling of ciprofloxacin analogues in complex with GyrA/GyrB**

The structure of *GyrA* and *GyrB* from *Arabidopsis thaliana* (Uniprot ID: Q9CAF6 and Q9SS38) was modelled with Alphafold 2.0 [PMID: 34265844], using the X-ray crystallography structure of the same proteins from *Staphylococcus aureus* (PDB ID: 2XCS, PMID: 20686482). The resulting computational model was then superimposed with the experimental model, thereby aligning the computational model with the dsDNA helix, which constitutes a large portion of the ciprofloxacin (CFX) binding site.

The computational model, along with the dsDNA, were then prepared using Protein Preparation Wizard (Schrodinger LLC, NY, USA) [PMID: 23579614]. Ciprofloxacin, **5** and **6** were prepared for docking using ligprep [PMID: 17899391]. Docking of these molecules was done with Glide Standard Precision mode [PMID: 15027865, 15027866], in two stages. Initially, ciprofloxacin was docked with no constraints, into the binding site. The highest scoring pose had ciprofloxacin sandwiched between the nitrogen bases of the dsDNA. This pose was used as core constraint, with maximum of 0.10Å RMSD tolerated between the core and the matching sections of **5** and **6**. This was necessary because of the long flexible tail added to the analogues, which makes exhaustively sampling all the different conformations computationally prohibitive and the unconstrained docking results were therefore poor in quality.

**Gyrase inhibition assay**

Gyrase inhibition assay was performed using TopoGEN *E. coli* DNA gyrase and relaxed DNA assay kit (TG2000G-1KIT) according to the manufacturer's instructions.

**Plants phenotypic characterization**

*Arabidopsis thaliana* seeds were sown and grown in pots as described in "Plant material and growth conditions" section. On the $10^{th}$ and $12^{th}$ day each pot was sprayed with 1 ml DDW containing 0.05% TWEEN 20 (Merk) and CFX (10 μM) or **6** (50 μM). Morphological and photosynthesis parameters were analyzed with the PlantScreen$^{TM}$ Phenotyping System, Photon Systems Instruments (PSI), Czech Republic. Plants were sown in PSI standard pots and imaged once a day from day 14 to day 26. On the last measuring day plants were also photographed using Nikon D5300 with AF-S micro Nikkor 60 mm lens.

#### Root elongation assay

*Arabidopsis thaliana* seedlings were grown on agar as described in "Plant material and growth conditions" section. On the 5$^{th}$ day the seedlings were transferred onto MS plates containing various concentrations of CFX or **6**. The seedlings were placed in growth chamber under long day condition and root length was marked daily. On the 10$^{th}$ day the plants were scanned and root length was measured using FIJI software.

#### Transmission electron microscopy (TEM) imaging

*Arabidopsis thaliana* plants were grown in pots as described in "Plant material and growth conditions" section. On the 10$^{th}$ and 12$^{th}$ day each pot was sprayed 1 ml DDW containing 0.05% TWEEN 20 (Merk) and CFX (10 μM) or **6** (50 μM). On the 14$^{th}$ day, approximately 1x1 cm$^2$ tissues were cut from leaves. Tissues were fixed in 2.5% glutaraldehyde in PBS over night at 4 °C. After several washings in PBS tissues were post fixed in 1% $OsO_4$ in PBS for 2 h at 4 °C. Dehydration was carried out in graded ethanol followed by embedding in glycid ether. Thin sections were mounted on Formvar/Carbon coated grids, stained with uranyl acetate and lead citrate and examined in Jeol 1400 – Plus transmission electron microscope (Jeol, Japan). Images were captured using SIS Megaview III and iTEM the Tem imaging platform (Olympus).

#### Relative DNA expression

*Arabidopsis thaliana* seedlings were grown on agar as described in "Plant material and growth conditions" section. On the 5$^{th}$ day seedlings were transferred onto MS plates containing CFX (10 µM) or **6** (50 µM) for 24 h. About 100 seedlings were harvested, frozen in liquid nitrogen, and crushed by a TissueLyser to form a thin powder. 500 µl phenol-chloroform (1:1) buffer was added to each sample, mixed well, and centrifuged for 3 min. The supernatant was transferred into a new tube with an equal volume of chloroform to remove phenol traces, centrifuged for 3 min, and the supernatant was transferred to a new tube. An equal volume of 5 M ammonium acetate was added to the supernatant to a final concentration of 2.5 M, followed by the addition of 2-2.5 ml of cold 100% EtOH. Then, samples were stored at -20° C for 12 h, and centrifuge at 14,000 rpm for 15 min. Supernatants were decanted. 1 ml of 70% EtOH was added to the pellet, and the mixture was centrifuged for 1 min. The supernatant was discarded, and the pellets were air-dried for 10 min and resuspended in TE buffer. Quantitative

RT–PCR was performed with 10 ng DNA in a final volume of 10 µl with Fast SYBR™ Green Master Mix (Applied Biosystems, Cat. No. 4385612) using StepOnePlus™ System and software (Thermo Fisher Scientific). The reaction conditions included 40 amplification cycles, (3 sec at 95 °C, 30 sec at 60 °C). Three technical repeats were performed for each DNA sample, and at least three biological repeats were used for each treatment. The relative quantification was calculated with the ΔΔCt method, TUB2 (TUBULIN BETA CHAIN 2) and ACT2 (ACTIN 2) were used as reference genes. Primers are specified in Supplementary Table S1.

**Table S1:** List of primers used for the relative DNA expression qRT-PCR.

| Gene | AGI | Sequence |
|---|---|---|
| TUB2 | AT5G62690 | F:TGGCATCAACTTTCATTGGA<br>R:ATGTTGCTCTCCGCTTCTGT |
| ACT2 | AT3G18780 | F:GAGATGGAGACCTCGAAAACC<br>R:TTGTCCGTCGGGTAATTCAT |
| PSBK | ATCG00070 | F: AGGCCTACGCCTTTTTAAT<br>R: GCTTGCCAAACAAAGGCTAA |
| YCF3 | AT5G44650 | F:ATGTCGGCTCAATCTGAAGG<br>R:TTCGTTCTAATGCCCGAAAA |
| COX2 | ATMG00160 | F:TTCCGATGAGCAGTCACTCA<br>R:TGAGGAAGGTACAGCCCAAC |
| RPS3 | AT3G07040 | F:CTCGCTCTTTCCATTCTTCG<br>R:ATCCTTACCCACGAGGTTCC |

**DCF staining**

*Nicotiana benthamiana* plants were grown as described in “Plant material and growth conditions” section. At about 2 weeks old leaves were infiltrated with **6** (50 µM) or DDW. After 3 h leaves were infiltrated again with MitoTracker red (1 µM) and 2',7'-dichlorodihydrofluorescein diacetate (DCF, 50 µM). After 10 min leaves were imaged. In each image, 5 mitochondria and 5 chloroplasts were chosen randomly and DCF emission was measured using ZEN 3.1 blue edition software.

**Relative RNA expression**

*Arabidopsis thaliana* seedlings were grown on agar as described in “Plant material and growth conditions” section. On the 5$^{th}$ day seedlings were transferred onto MS plates containing CFX (10 µM) or **6** (50 µM) or mock treatment. After 3 or 6 h seedlings were harvested and grounded in liquid nitrogen. Total RNA was extracted from 70 to 100 mg grounded plant tissue using RNeasy Plant Mini Kit (QIAGEN, Cat. No. 74904), according to the manufacture protocol. DNA was removed by DNase I, RNase-free (Thermo Fisher Scientific, Cat. No.  EN0521). Total RNA (2 µg) was converted to complementary DNA (cDNA) using High-Capacity cDNA Reverse Transcription Kit (applied biosystems by Thermo Fisher Scientific, Cat. No. 4368814) according to manufacturer protocols. Quantitative RT–PCR was performed with 40 ng cDNA in a final volume of 20 µl, with Fast SYBR™ Green Master Mix (Applied Biosystems, Cat. No. 4385612) using the QuantStudio1 Real-Time PCR System (Thermo Fisher Scientific). The reaction conditions included 40 amplification cycles (3 sec at 95°C, 30 sec at 60 °C). Three technical repeats were performed for each cDNA sample, and three biological repeats were used for each treatment. Relative quantification was calculated using the ΔΔCt method, with PP2AA3 (PROTEIN PHOSPHATASE 2A SUBUNIT A3) used as the reference gene. Primers are specified in Supplementary Table S2.

**Table S2:** List of primers used for the relative RNA expression qRT-PCR.

| Gene | AGI | Sequence |
|---|---|---|
| AOX1A | AT3G22370 | F:CCATGGGAAACGTATAAAGCTGAT<br>R:CCATATCTCCTCTGGAAGAACA |
| ANAC013 | AT1G32870 | F:GGAATTACCTGAGAAAGCGTTGTA<br>R:TGCTTTCCAATAGCCACGTT |
| PP2AA3 | AT1G13320 | F: TAACGTGGCCAAAATGATGC<br>R: GTTCTCCACAACCGCTTGGT |

**Data analysis**

All graphs (line charts, bar charts, box plots) were done using OriginPro 2024. Statistical analysis were done either with Microsoft Excel (*t*-test) or OriginPro2024 (ANOVA). Co-localization analysis were done using FIJI. Grey value measures were done either with FIJI (fluorescent probes imaging) or ZEN 3.1 blue edition (DCF staining). $EC_{50}$ calculations (DoseResp fitting curve) were done using OriginPro2024. Figures were made using Photoshop.

### Chemical methods

#### General

All chemicals were purchased from Sigma-Aldrich, TCI or CombiBlocks and used as received unless otherwise stated. Anhydrous solvents and reagents (THF, DMF, and toluene) were obtained as SureSeal bottles from Sigma-Aldrich. Thin-layer chromatography and flash chromatography were performed using Merck KGaA pre-coated silica gel 60 F-254 plates and Silicycle silica gel 40-63 (230-400 mesh), respectively. UV absorbance spectra were recorded on Agilent Cary 60 UV-Vis Spectrophotometer. Fluorescence spectra were recorded on Fluorolog 2 (Spex) fluorimeter. Low resolution ESI mass spectrometry was performed on LC/MS Acquity QDa detector coupled with Waters HPLC. High resolution ESI mass spectrometry was performed on a Waters SYNAPT system. $^{1}H$ and $^{13}C$ NMR spectra were collected in DMSO-d6 (Cambridge Isotope Laboratories, Cambridge, MA) at 25 $^{o}C$ using a Bruker Advance III spectrometer at 400 MHz and 100 MHz, respectively, at the Department of Chemistry NMR Facility at Tel-Aviv University. All chemical shifts are reported in the standard δ notation of parts per million using the either TMS or residual solvent peak as an internal reference. Abbreviations: $PPh_3$ - triphenylphosphine; MeCN - acetonitrile; DCM - dichloromethane; DMF - dimethylformamide; THF - tetrahydrofuran; MeOH - methanol; EtOH - ethanol; NBD-Cl - 4-chloro-7-nitrobenzofurazan; TAMRA-SE - 5-(and-6)-carboxytetramethylrhodamine succinimidyl ester, $Et_3N$ - trimethylamine; DMAP - dimethylaminopyridine; DIPEA - *N*,*N*-Diisopropylethylamine; HBTU - (2-(1H-benzotriazol-1-yl)-1,1,3,3-tetramethyluronium hexafluorophosphate; TFA - trifluoroacetic acid; RT - room temperature.

#### LC/MS Analysis conditions

HPLC-MS analysis was performed on Waters HPLC with XBridge C18 column (100 X 3 mm, 5 µm) using water (solvent A) and acetonitrile (solvent B), both containing 0.1% TFA as an additive at flow rate of 1 mL/min. The program consists of 1 min 100% A, then 15 min gradient from 0% to 100% B, following by 1 min 100% B, then 2 min gradient from 0% to 100% A and finally 1 min at 100% A. Mass spectrometry was performed on LC/MS Acquity QDa detector coupled with Waters HPLC.

#### Preparative HPLC purification conditions

Preparative HPLC was performed on Waters 2545 HPLC with XBridge C18 column (100 X 19 mm, 5 µm) using water (solvent A) and acetonitrile (solvent B), both containing 0.1% TFA as an additive at flow rate of 15 mL/min. The program consists of 2 min 100% A, then 22 min gradient from 0% to 100% B, following by 2 min 100% B, 2 min gradient from 0% to 100% A and finally 2 min 100% A.

### Synthetic procedures

#### Synthesis of 8

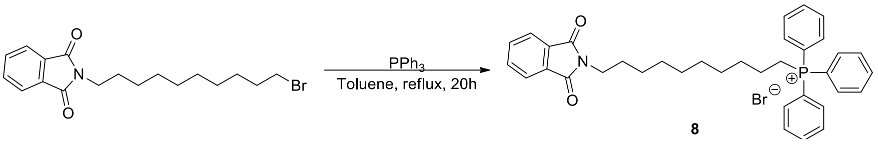

**Scheme S1:** Synthetic scheme of **8**.

**8** was synthesized according to an existing protocol.[6] Briefly, triphenylphosphine (120 mg, 0.46 mmol) and bromophtalimide (252 mg 0.69 mmol) were dissolved in dry toluene (3 mL) and the mixture was stirred and refluxed under argon atmosphere. After 20 h, the solvent was removed under reduced pressure and the crude residue was purified by flash column chromatography (DCM:EtOH 80:20) to yield **8** as a yellow oil (292 mg, 46%). $^{1}$H NMR (400 MHz, DMSO) δ 8.11 – 7.89 (m, 19H), 3.84 – 3.74 (m, 2H), 2.45 (t, J = 7.4 Hz, 3H), 1.80 – 1.53 (m, 10H), 1.39 – 1.30 (m, 6H). $^{13}$C NMR (101 MHz, DMSO) δ (ppm): 172.9, 167.9, 134.9, 134.4, 133.6, 133.5, 131.6, 130.3, 130.2, 123.0, 119.0, 118.2, 37.4, 31.3, 29.0, 28.9, 28.7, 28.4, 27.8, 26.2, 24.4, 22.1.. HR-MS (ESI) calcd. for formula $C_{36}H_{39}NO_2P$ $[M-Br]^+$: 548.2718; found: 548.2721.

#### Synthesis of 9

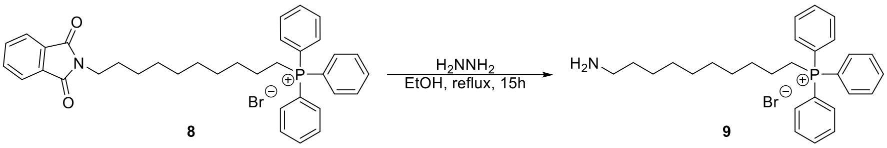

**Scheme S2:** Synthetic scheme of **9**.

**9** was synthesized according to an existing protocol.[6] Briefly, to a solution of **8** (252 mg, 0.45 mmol) in EtOH (1.5 mL) was added hydrazine (22.5 µL, 0.45 mmol) and the mixture was stirred under reflux. After 15 h, the solvent was removed under reduced pressure, and the crude residue was first purified by flash column chromatography (DCM:EtOH 80:20). The product was then pass through preparative HPLC, using the conditions listed above to yield **9** as a yellow oil (106 mg, 47%). $^{1}$H NMR (400 MHz, DMSO) δ $^{1}$H NMR (400 MHz, DMSO) δ 8.00 – 7.72 (m, 15H), 3.62 (dd, *J* = 5.9, 4.3 Hz, 2H), 2.75 (q, *J* = 6.5 Hz, 2H), 1.61 – 1.34 (m, 10H), 1.20 (s, 6H). $^{13}$C NMR (101 MHz, DMSO) δ (ppm): 134.9, 133.6, 133.5, 130.3, 130.2, 119.0, 118.1, 29.8, 29.0, 28.9, 28.7, 28.5, 28.2, 27.0, 25.8, 22.1, 14.0. HR-MS (ESI) calcd. for formula $C_{28}H_{37}NP$ $[M-Br]^+$: 418.2664; found: 418.2660.

**Synthesis of 1**

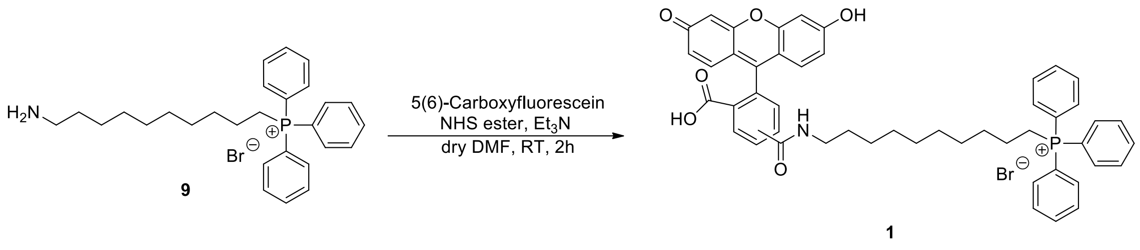

**Scheme S3:** Synthetic scheme of **1**.

**1** was synthesized using a modified protocol.[7] 5(6)-Carboxyfluorescein NHS ester (5 mg, 10 µmol) was dissolved in dry DMF (1 ml) then **9** (5 mg, 12 µmol) was added, followed by addition of trimethylamine (1.66 µL, 12 µmol). The reaction was stirred for 2 h at room temperature and the product was directly purified via preparative HPLC using the conditions listed above to yield **1** as a red-orange powder (11 mg, 90%). $^{1}$H NMR (400 MHz, DMSO) δ (ppm): 10.18 (s, 4H), 8.79 (t, J = 5.6 Hz, 1H), 8.65 (t, J = 5.6 Hz, 1H), 8.45 – 8.43 (m, 1H), 8.22 (dd, J = 8.0, 1.6 Hz, 1H), 8.15 (dd, J = 8.1, 1.4 Hz, 1H), 8.07 (d, J = 8.1 Hz, 1H), 7.93 – 7.85 (m, 6H), 7.84 – 7.73 (m, 24H), 7.65 (t, J = 1.0 Hz, 1H), 7.36 (d, J = 8.1 Hz, 1H), 6.71 – 6.67 (m, 4H), 6.60 – 6.51 (m, 8H), 3.59 – 3.49 (m, 4H), 3.29 (q, J = 6.7 Hz, 2H), 3.17 (q, J = 6.7 Hz, 2H), 1.58 – 1.38 (m, 12H), 1.37 – 1.09 (m, 22H). $^{13}$C NMR (101 MHz, DMSO) δ (ppm): 168.2, 168.0, 164.4, 164.3, 159.6, 158.0, 157.6, 154.6, 151.8, 140.8, 136.4, 134.9, 134.8, 134.6, 133.6, 133.5, 130.3, 130.1, 129.3, 129.2, 129.1, 128.1, 126.4, 124.8, 124.2, 123.1, 122.1, 119.0, 118.1, 112.7, 112.6, 109.1, 109.1, 102.3, 102.2, 29.9, 29.8, 29.7, 29.6, 28.9, 28.8, 28.7, 28.6, 28.0, 27.8, 26.4, 26.4, 21.7, 21.7, 21.6, 20.4, 19.9. HR-MS (ESI) calculated $C_{49}H_{47}NO_6P$ [M-Br]$^{+}$: 776.3136; found: 776.3136.

**Synthesis of 2**

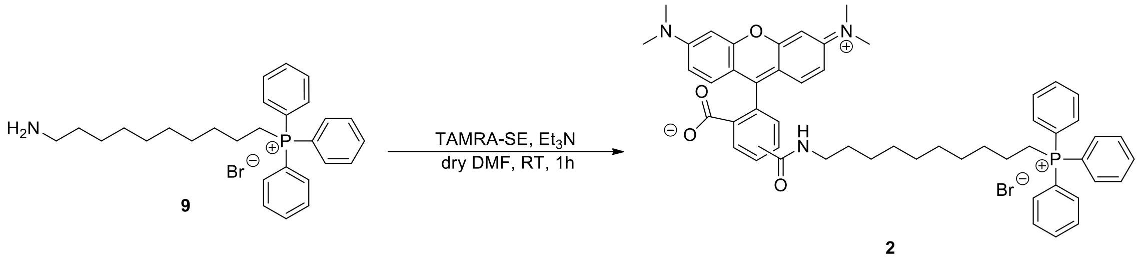

**Scheme S4:** Synthetic scheme of **2**.

**2** was synthesized using a modified protocol.[7] TAMRA-SE (4.5 mg, 8.6 µmol) dissolved in dry DMF (1 ml), then **9** (4 mg, 9.5 µmol) was added, followed by addition of triethylamine until reaching pH ≈ 9 (8 µL). The reaction was stirred for 1 h at room temperature and the product was directly purified via preparative HPLC using conditions listed above to yield **2** as a red-purple powder (4.7 mg, 60%).

$^{1}H$ NMR (400 MHz, DMSO) δ (ppm): 8.86 (t, J = 5.6 Hz, 1H), 8.74 (t, J = 5.6 Hz, 1H), 8.66 (s, 1H), 8.28 (dd, J = 7.9, 1.7 Hz, 2H), 8.22 (dd, J = 8.2, 1.8 Hz, 1H), 7.92 – 7.74 (m, 32H), 7.56 (d, J = 8.0 Hz, 1H), 7.08 – 6.92 (m, 11H), 3.58 – 3.54 (m, 4H), 3.32 (q, J = 6.7 Hz, 4H), 3.25 (s, 24H), 1.57 – 1.41 (m, 12H), 1.32 – 1.18 (m, 20H). $^{13}C$ NMR (101 MHz, DMSO-d6) δ (ppm): 164.5, 164.2, 157.9, 157.6, 136.2, 134.9, 134.8, 133.6, 133.5, 130.3, 130.1, 128.9, 119.0, 118.1, 96.3, 40.5, 29.9, 29.7, 29.0, 28.9, 28.8, 28.8, 28.7, 28.6, 28.1, 28.0, 26.5, 21.7, 20.4, 19.9. HR-MS (ESI) calculated $C_{53}H_{57}NO_4P$ [M-Br]$^+$: 830.4087; found: 830.4084.

**Synthesis of 3**

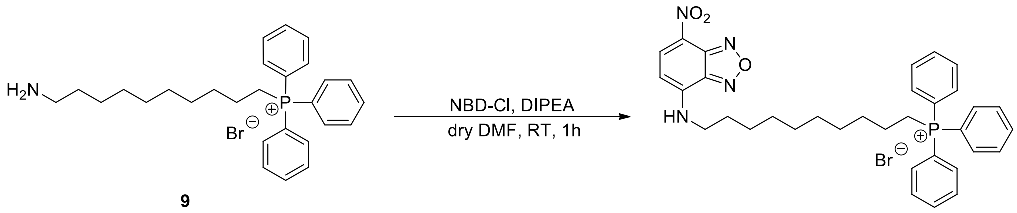

9

3

**Scheme S5:** Synthetic scheme of **3**.

**3** was synthesized using a modified protocol.[8] NBD-Cl (8 mg, 40 µmol) was dissolved in dry DMF (2 ml) with DIPEA (14.5 µL, 80 µmol), then **9** (20 mg, 40 µmol) was added. The reaction was stirred for 1 h at room temperature and the product was directly purified via preparative HPLC using conditions listed above to yield **3** as a red-orange powder (22 mg, 83%). $^{1}H$ NMR (400 MHz, DMSO) δ (ppm): 9.54 (t, J = 6.0 Hz, 1H), 8.51 (d, J = 8.9 Hz, 1H), 7.92 – 7.74 (m, 15H), 6.40 (d, J = 9.0 Hz, 1H), 3.48 – 3.41 (m, 2H), 1.64 (q, J = 7.3 Hz, 2H), 1.54 – 1.39 (m, 4H), 1.37 – 1.15 (m, 12H). $^{13}C$ NMR (101 MHz, DMSO) δ (ppm): 145.6, 144.9, 144.7, 138.4, 135.4, 135.3, 134.1, 134.0, 130.8, 130.6, 121.0, 119.5, 118.6, 99.5, 43.8, 30.4, 30.2, 29.3, 29.1, 28.6, 28.1, 26.8, 22.2, 22.2, 20.8, 20.3. HR-MS (ESI) calculated $C_{34}H_{38}N_4O_3P$ [M-Br]$^+$: 581.2682; found: 581.2685.

**Synthesis of 10**

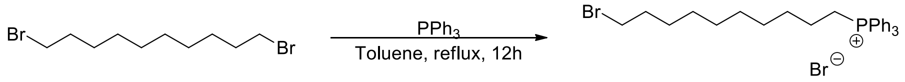

10

**Scheme S6:** Synthetic scheme of **10**.

**10** was synthesized according to an existing protocol.[9] Briefly, triphenylphosphine (291 mg, 1.1 mmol) and 1,10-dibromodecane (500 mg 1.6 mmol) were dissolved in dry toluene (3 mL) and the mixture was refluxed under argon atmosphere. After 12h, solvent was removed under reduced pressure and the crude was purified by flash column chromatography (DCM:MeOH 95:5) to yield **10** as a brown oil (286 mg, 46%).$^{1}$H NMR (400 MHz, DMSO) δ (ppm): 8.00 – 7.72 (m, 15H), 3.35 (t, J = 6.5 Hz, 2H), 1.53 – 1.33 (m, 6H), 1.30 – 1.11 (m, 12H). $^{13}$C NMR (101 MHz, DMSO) δ (ppm): 134.9, 133.6, 133.5, 130.9, 130.2, 119.0, 118.2, 35.2, 32.9, 29.8, 29.7, 28.7, 28.5, 28.0, 27.5, 21.7, 20.4, 19.9. LC/MS: Retention time 13.83 min, 481.26 $[M-Br]^+$.

**Synthesis of 4**

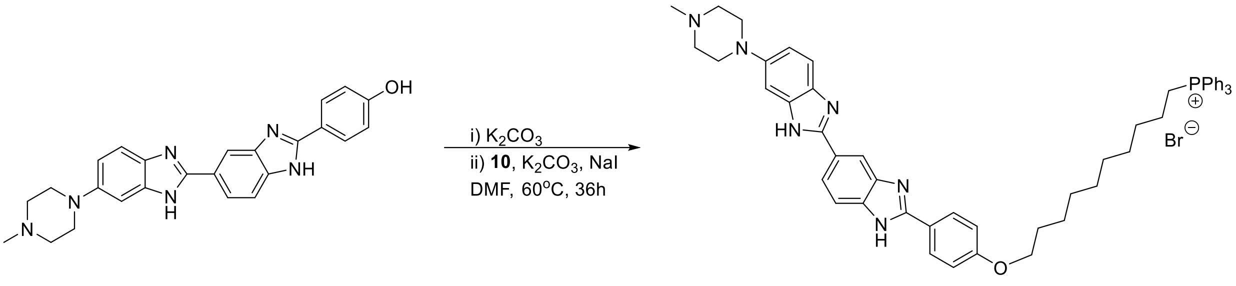

**4**

**Scheme S7:** Synthetic scheme of **4**.

Hoechst dye 33258 (30 mg, 48 µmol) and **10** (34 mg, 72 µmol) were dissolved in DMF (1 ml) and potassium carbonate (18 mg, 130 µmol) was added. The reaction was heated to 60 °C and left to stir. After 12 h, **10** (46 mg, 96 µmol) and potassium carbonate (20 mg, 144 µmol) were added again, along with a catalytic amount of sodium iodide. After 24 h, the solvent was removed under reduced pressure, and the crude was purified by flash column chromatography (DCM:MeOH 80:20) and purified again via preparative HPLC using the conditions listed above to yield **4** as a red powder (30 mg, 77%). $^{1}$H NMR (400 MHz, DMSO-d6) δ (ppm): 8.40 – 8.22 (m, 1H), 8.16 (dd, J = 8.7, 5.2 Hz, 2H), 8.01 (dd, J = 21.6, 8.4 Hz, 1H), 7.94 – 7.64 (m, 15H), 7.57 – 7.43 (m, 3H), 7.13 – 7.08 (m, 2H), 6.93 (d, J = 9.0 Hz, 1H), 4.08 – 4.01 (m, 2H), 3.62 – 3.50 (m, 2H), 3.16 (s, 4H), 2.61 (s, 4H), 2.32 (s, 3H), 1.77 – 1.68 (m, 2H), 1.53 – 1.20 (m, 16H). $^{13}$C NMR (101 MHz, DMSO-d6) δ (ppm): 134.9, 134.8, 133.6, 133.5, 130.4, 130.3, 130.6, 130.2, 130.1, 130.1, 128.6, 128.5, 119.0, 118.1, 114.8, 67.6, 54.6, 28.8, 28.7, 28.6, 28.4, 25.4. HR-MS (ESI) calculated $C_{53}H_{58}N_6OP$ $[M-Br]^+$: 825.4410; found: 825.4407.

**Synthesis of 5**

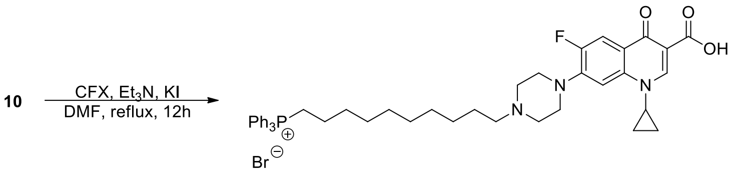

**Scheme S8:** Synthetic scheme of **5**.

**5** was synthesized using a modified protocol.[10] To a solution of CFX (47 mg, 0.14 mmol) in DMF (2 ml) was added **10** (104 mg, 0.21 mmol), followed by the addition of triethylamine (40 µl, 0.28 mmol) and potassium iodide (35 mg, 0.21 mmol). After 12 h, solvent was evaporated under reduced pressure and the crude was subjected to flash column chromatography (DCM:MeOH 80:20) then purified via preparative HPLC using conditions listed above to yield **5** as a yellow powder (68 mg, 60%). $^{1}$H NMR (400 MHz, DMSO-d6) δ (ppm): 8.63 (s, 1H), 7.92 – 7.87 (m, 3H), 7.84 – 7.74 (m, 13H), 7.55 (d, J = 7.4 Hz, 1H), 3.85 (tt, J = 7.5, 4.1 Hz, 1H), 3.64 – 3.55 (m, 2H), 2.90 (q, J = 7.3 Hz, 6H), 2.64 (s, 2H), 2.45 – 2.30 (m, 2H), 1.66 – 1.36 (m, 8H), 1.35 – 1.21 (m, 12H). $^{13}$C NMR (101 MHz, DMSO-d6) δ (ppm): 176.3, 176.3, 165.9, 154.2, 151.7, 147.9, 139.1, 134.9, 134.8, 133.6, 133.5, 130.3, 130.2, 119.0, 118.2, 111.0, 106.7, 106.4, 48.6, 41.3, 35.9, 29.9, 29.7, 28.9, 28.7, 28.1, 26.8, 21.8, 21.7, 20.5, 19.9, 11.0, 7.6. HR-MS (ESI) calculated $C_{45}H_{52}FN_3O_3P$ [M-Br]$^+$:732.3730; found: 732.3729.

**Synthesis of 11**

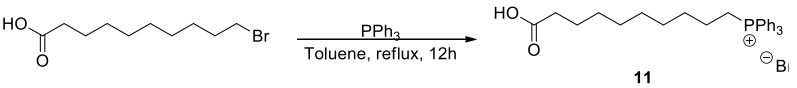

**Scheme S9:** Synthetic scheme of **11**.

**11** was synthesized according to an existing protocol.[9] Briefly, triphenylphosphine (696 mg, 2.6 mmol) and 10-bromodecanoic acid (1 gr 3.9 mmol) were dissolved in dry toluene (12 mL) and the mixture was refluxed under argon atmosphere. After 12 h, solvent was removed under reduced pressure, and the crude was purified by flash column chromatography (DCM:MeOH 90:10) to yield **11** as a white powder (534 mg, 40%). $^{1}$H NMR (400 MHz, DMSO) δ (ppm): 7.97 – 7.71 (m, 15H), 2.17 (t, *J* = 7.4 Hz, 2H), 1.63 – 1.12 (m, 16H). $^{13}$C NMR (101 MHz, DMSO) δ (ppm): 174.6, 134.9, 134.9, 133.7, 133.6, 130.3, 130.22, 119.0, 118.2, 33.7, 29.9, 29.7, 28.6, 28.6, 28.5, 28.0, 24.5, 21.7, 21.7, 20.4, 19.9. LC/MS: Retention time 13.67 min, 433.37 [M-Br]$^+$. HR-MS (ESI) calculated $C_{28}H_{34}O_2P$ [M-Br]$^+$: 433.2296; found: 433.2298.

**Synthesis of 6**

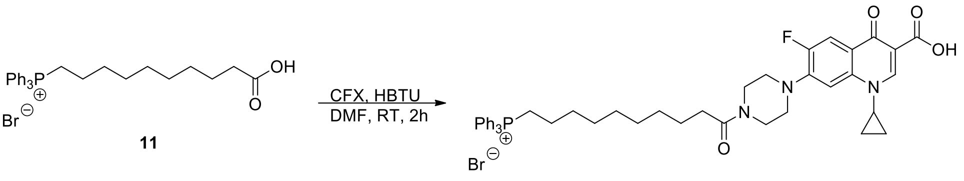

6

**Scheme S10:** Synthetic scheme of **6**.

To a solution of ciprofloxacine (CFX) (123 mg, 0.37 mmol) in DMF (2 ml) was added **11** (147 mg, 0.34 mmol), followed by the addition of HBTU (141 mg, 0.37 mmol) and DIPEA (65 µL, 0.32 mmol). After 2 h, the reaction was diluted with DCM (22 ml) and washed three times with 1 M HCl until the pH of the solution was ~5). The organic phase was dried over magnesium sulfate and DCM was removed under reduced pressure. The crude was purified by flash chromatography (DCM:MeOH 80:20) and solvents were removed under reduced pressure to yield **6** as a yellow powder (164 mg, 58%). $^{1}$H NMR (400 MHz, DMSO-d6) δ (ppm): 8.68 (s, 1H), 7.96 – 7.87 (m, 4H), 7.84 – 7.74 (m, 12H), 7.57 (d, J = 7.2 Hz, 1H), 3.85 – 3.78 (m, 1H), 3.67 (t, J = 5.0 Hz, 4H), 3.59 – 3.50 (m, 2H), 2.69 (s, 2H), 2.34 (t, J = 7.4 Hz, 2H), 1.58 – 1.39 (m, 6H), 1.33 – 1.17 (m, 14H). $^{13}$C NMR (101 MHz, DMSO-d6) δ (ppm): 176.4, 165.9, 154.1, 151.7, 148.1, 144.9, 144.8, 139.1, 138.4, 134.9, 134.8, 133.6, 133.5, 130.3, 130.1, 119.0, 118.8, 118.8, 118.1, 111.1, 110.9, 106.8, 106.6, 106.6, 49.7, 49.2, 44.5, 38.2, 35.8, 32.2, 30.9, 29.9, 29.7, 28.7, 28.0, 24.7, 22.0, 21.7, 20.4, 19.9, 13.9, 7.6. HR-MS (ESI) calculated $C_{45}H_{50}FN_3O_4P$ $[M-Br]^+$: 746.3523; found: 746.3524.

**Synthesis of 7**

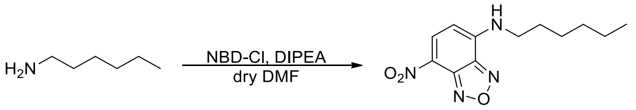

7

**Scheme S11:** Synthetic scheme of **7**.

**7** was synthesized according to a modified protocol.[8] NBD-Cl (50 mg, 0.25 mmol) was dissolved in dry DMF (5 ml) with DIPEA (87 µL, 0.5 mmol) and then hexylamine (33 µL, 0.25 mmol) was added. The reaction was stirred for 1 h in room temperature and directly purified via preparative HPLC using the conditions listed above to yield **7** as a red-orange powder (30.6 mg, 46%). $^{1}$H NMR (400 MHz, DMSO) δ (ppm): 9.54 (d, *J* = 5.6 Hz, 1H), 8.49 (d, *J* = 8.9 Hz, 1H), 6.39 (d, *J* = 9.0 Hz, 1H), 3.54 – 3.37 (m, 2H), 1.66 (h, *J* = 6.8 Hz, 2H), 1.41 – 1.24 (m, 6H), 0.90 – 0.82 (m, 3H). $^{13}$C NMR (101 MHz,

DMSO) δ (ppm): 145.1, 144.4, 144.2, 137.9, 120.5, 99.1, 43.4, 30.9, 27.6, 26.1, 22.0, 13.9. LC/MS: Retention time min, 265.12 $[M+H]^+$.

### $^{1}$H- and $^{13}$C-NMR spectra

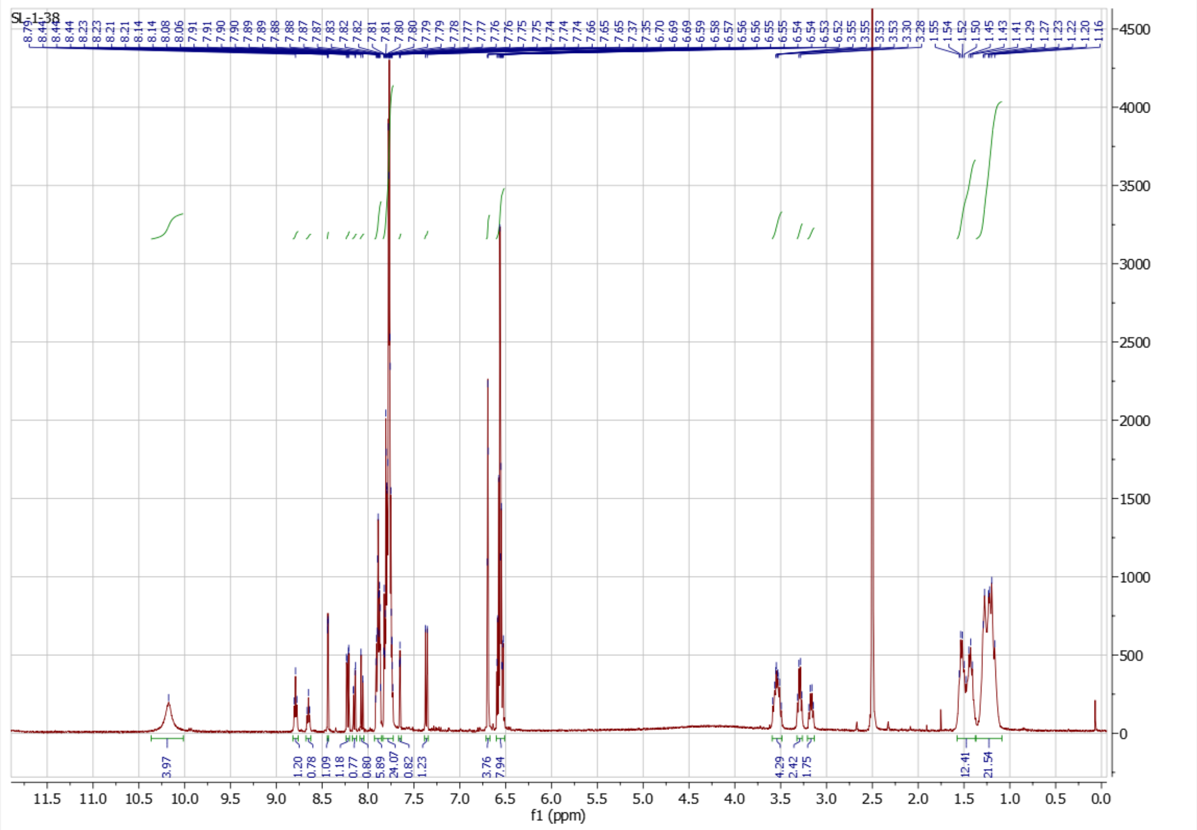

**Figure S5:** $^{1}$H NMR of **1** ON 400 MHz in DMSO-$d_6$.

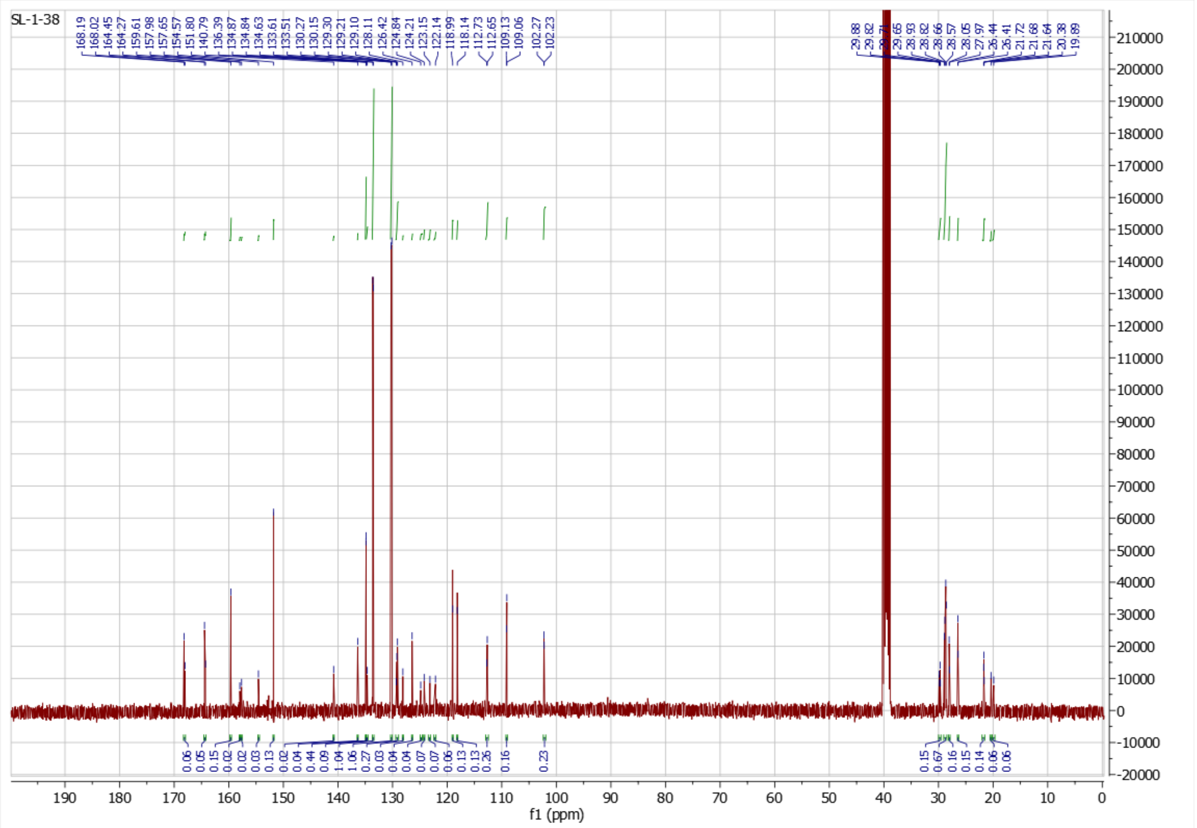

**Figure S6:** $^{13}$C NMR of **1** ON 101 MHz in DMSO-$d_6$.

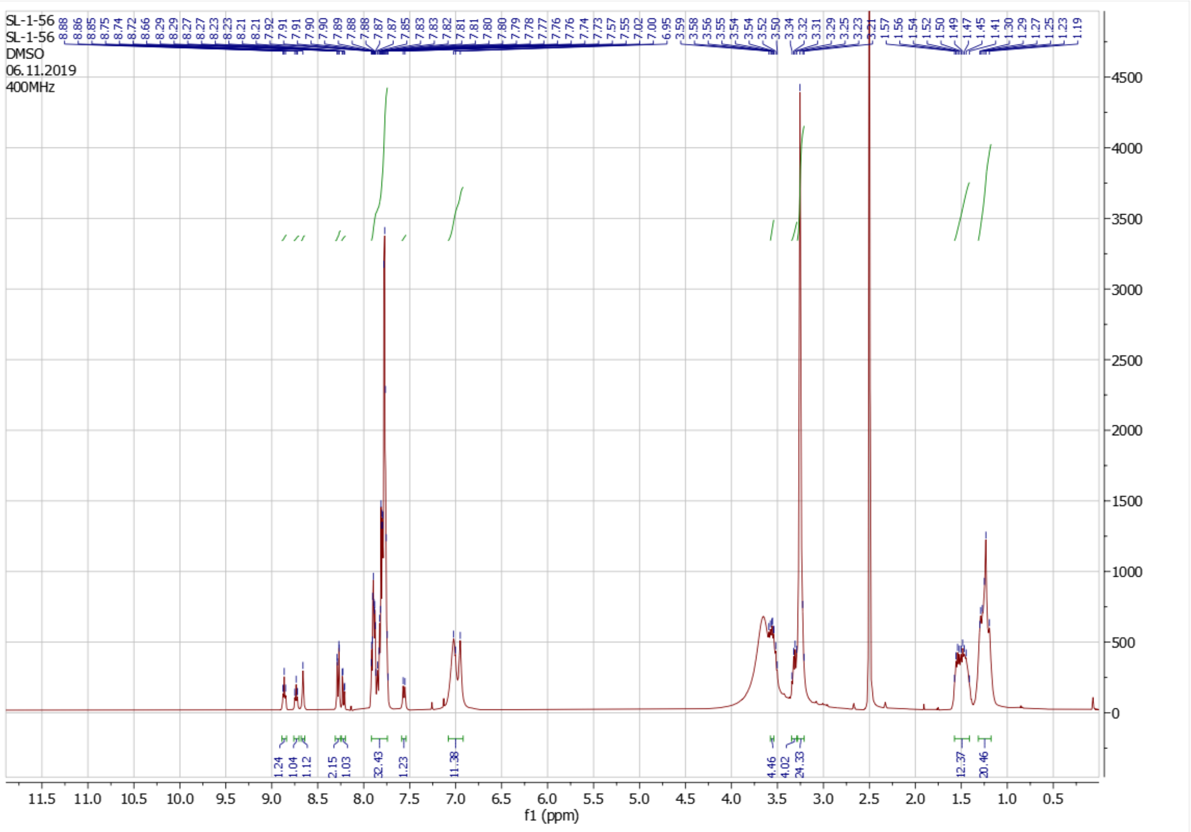

**Figure S7:** $^{1}$H NMR of **2** ON 400 MHz in DMSO-$d_6$.

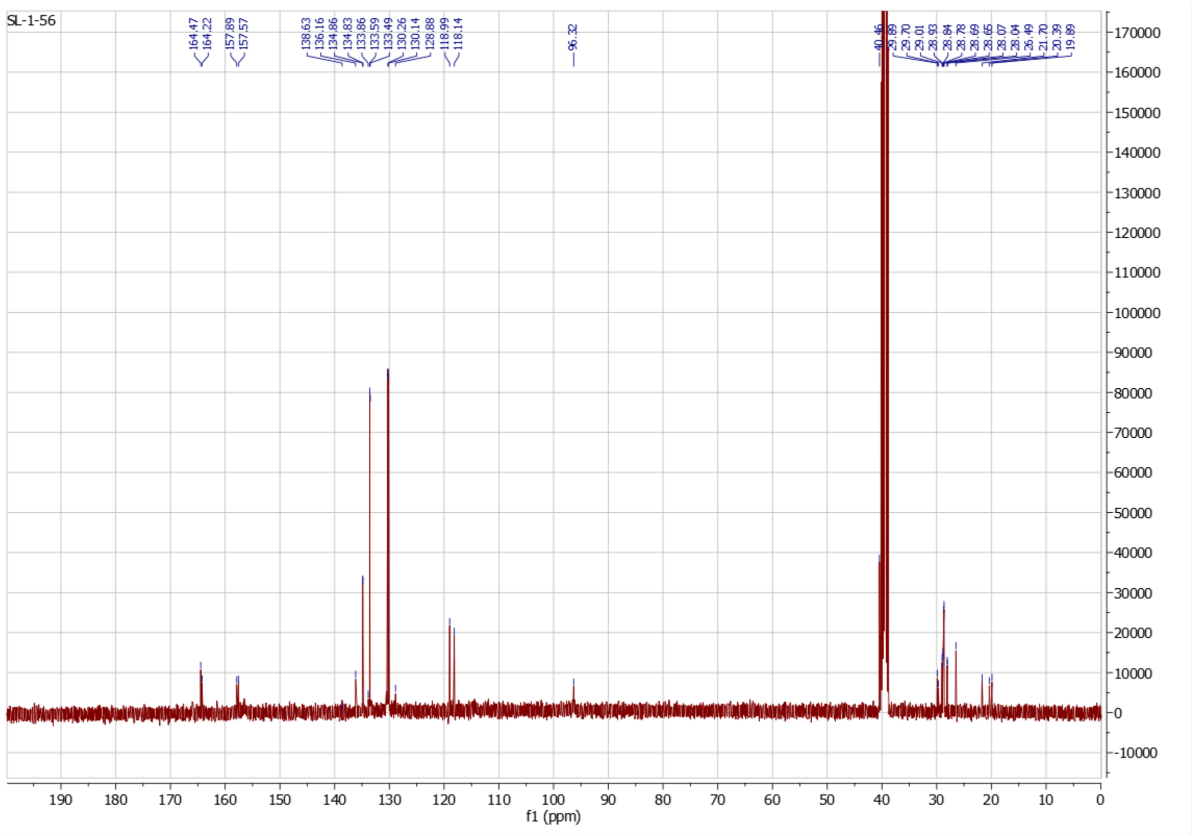

**Figure S8:** $^{13}$C NMR of **2** ON 101 MHz in DMSO-$d_6$.

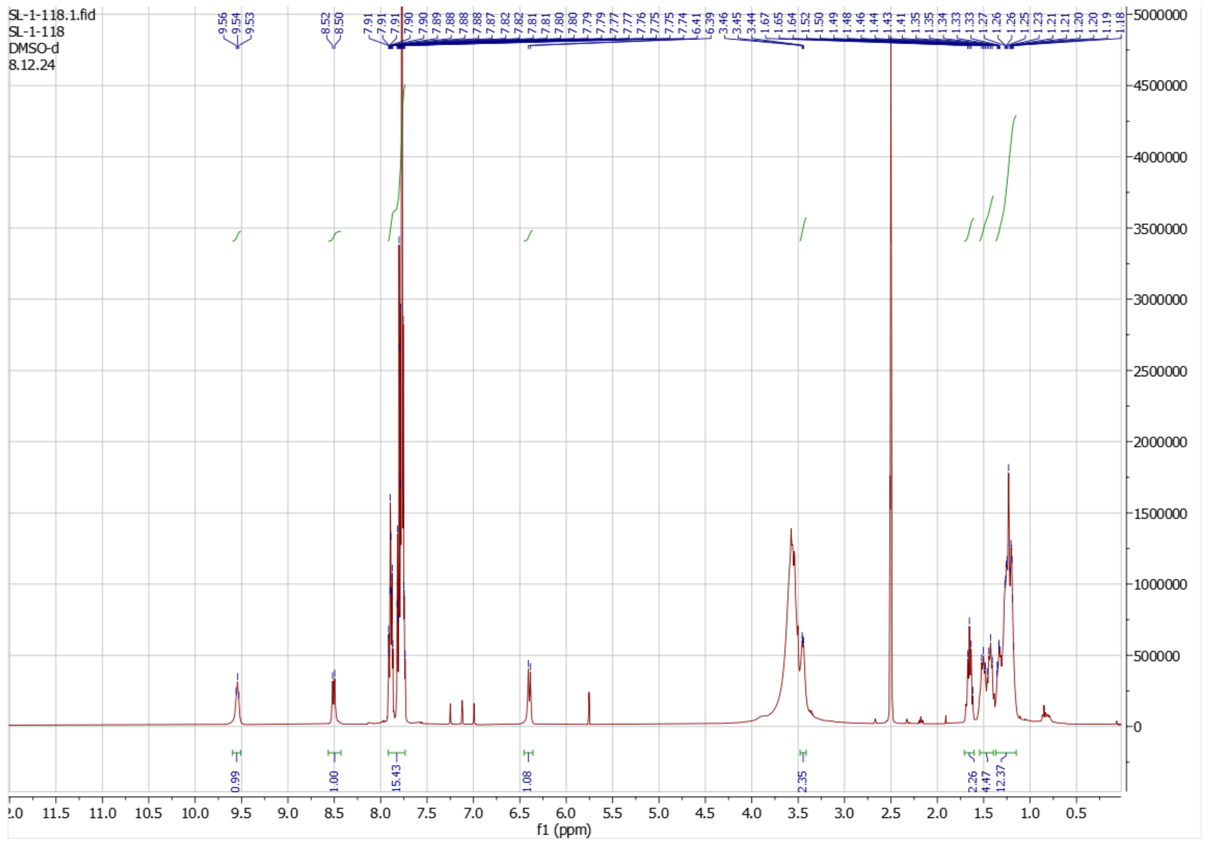

**Figure S9:** $^{1}$H NMR of **3** ON 400 MHz in DMSO-$d_6$.

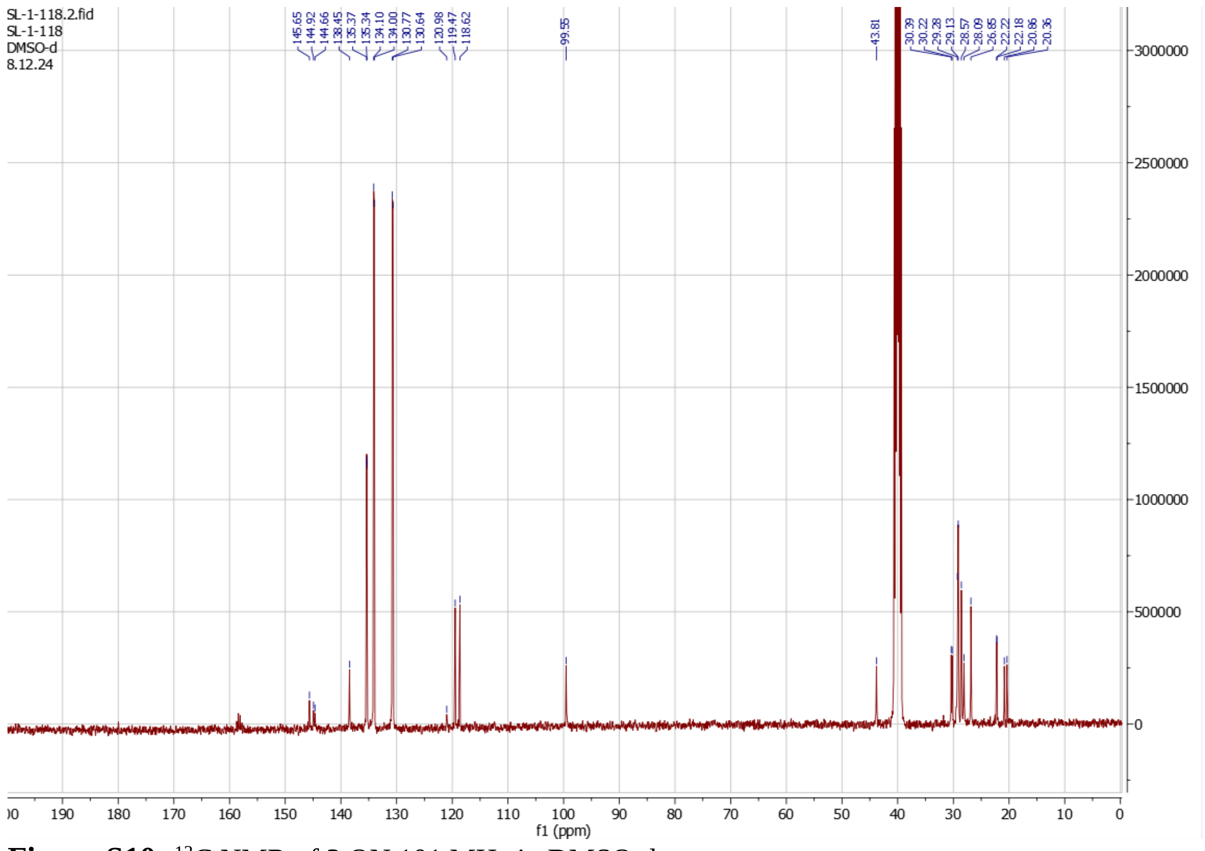

**Figure S10:** $^{13}$C NMR of **3** ON 101 MHz in DMSO-$d_6$.

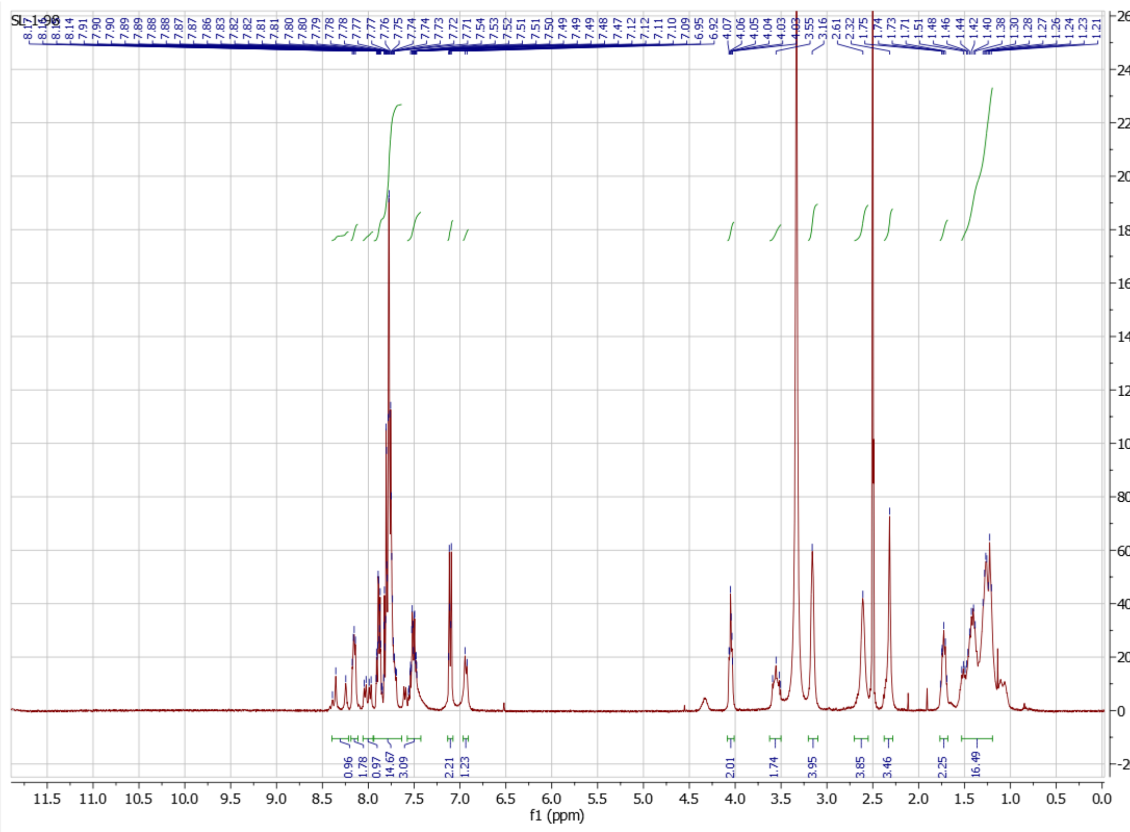

**Figure S11:** $^{1}$H NMR of **4** ON 400 MHz in DMSO-$d_6$.

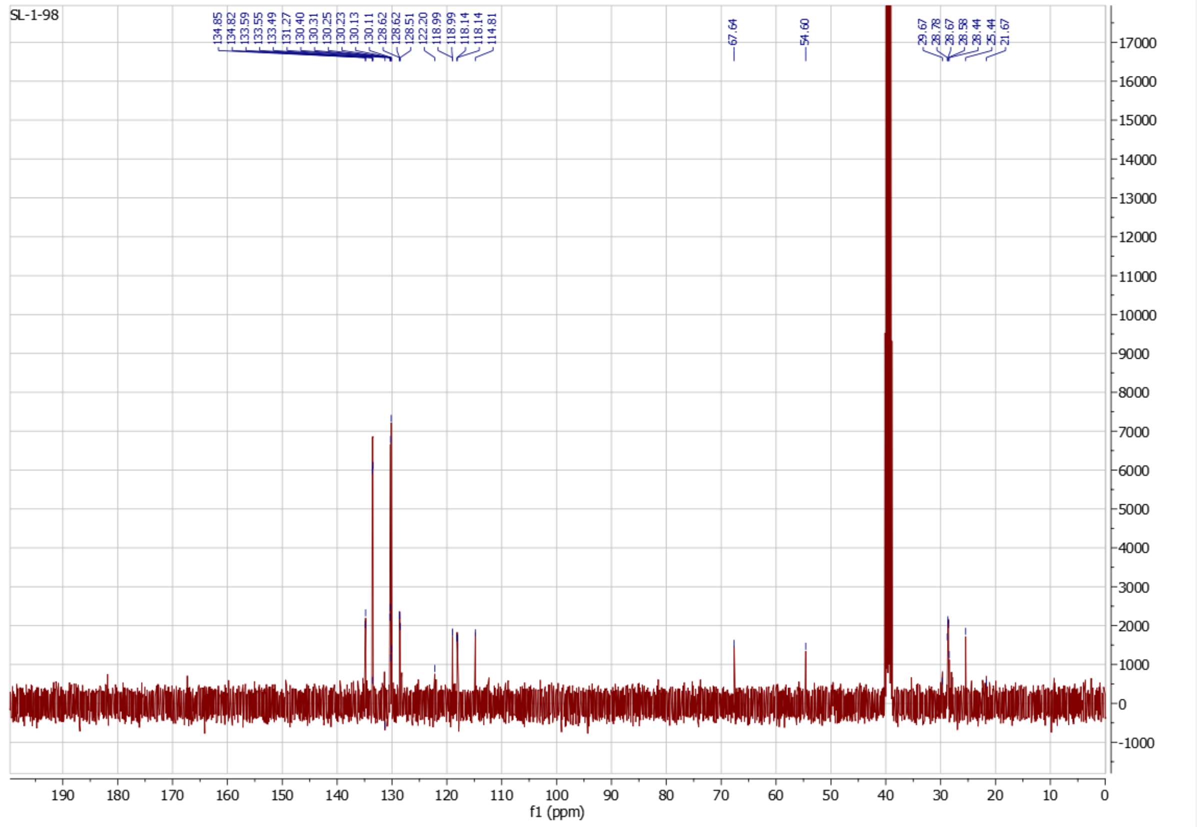

**Figure S12:** $^{13}$C NMR of **4** ON 101 MHz in DMSO-$d_6$.

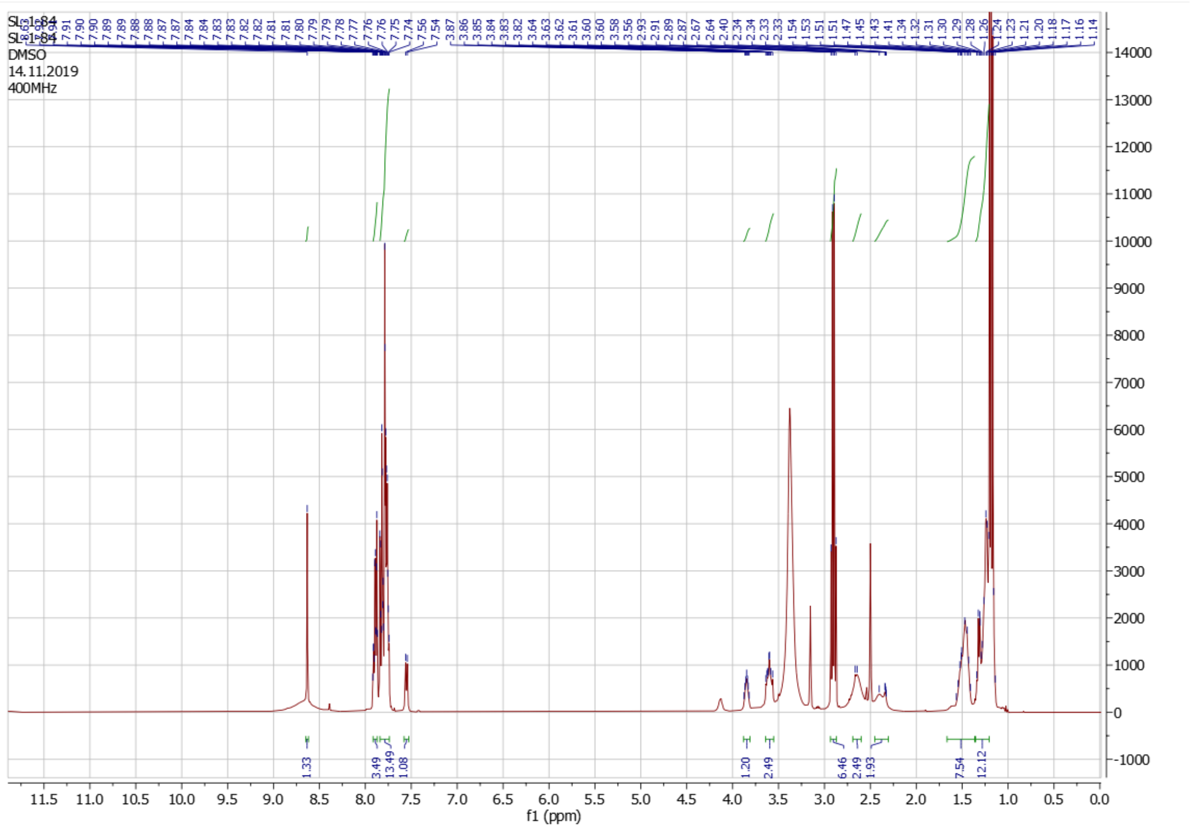

**Figure S13:** $^{1}$H NMR of **5** ON 400 MHz in DMSO-$d_6$.

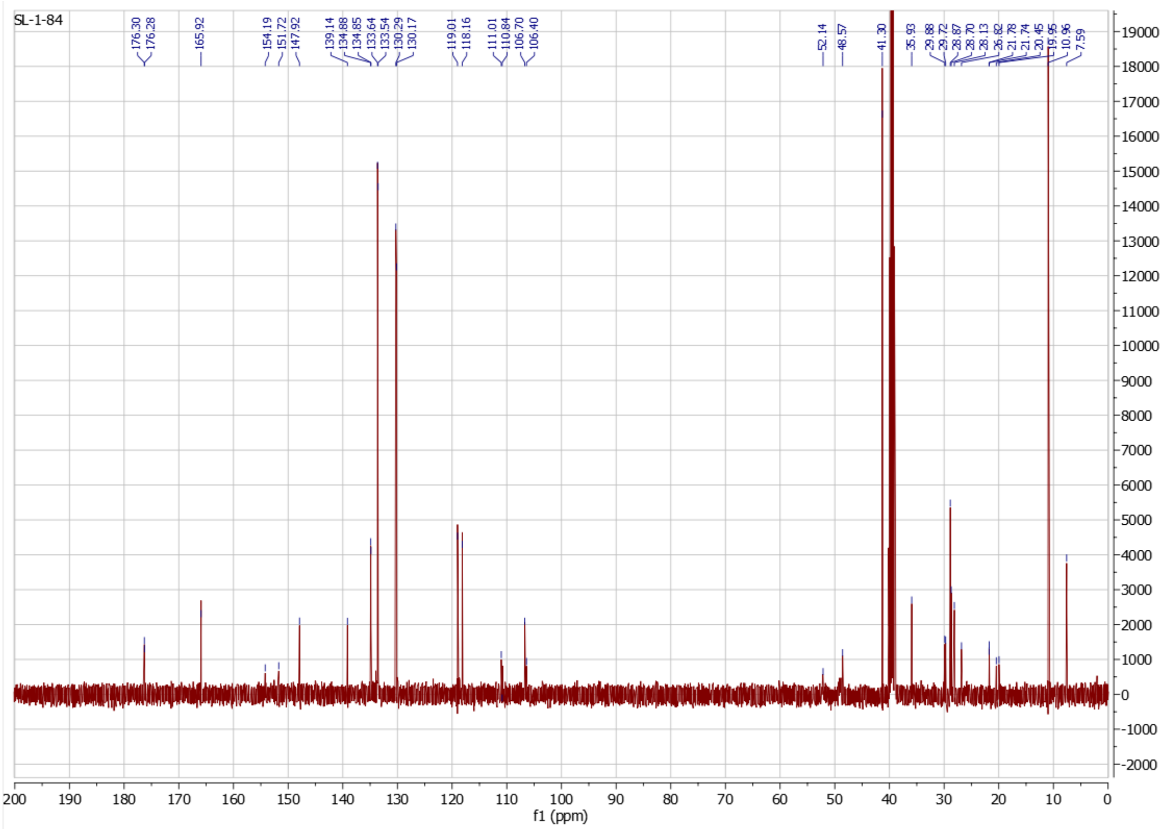

**Figure S14:** $^{13}$C NMR of **5** ON 101 MHz in DMSO-$d_6$.

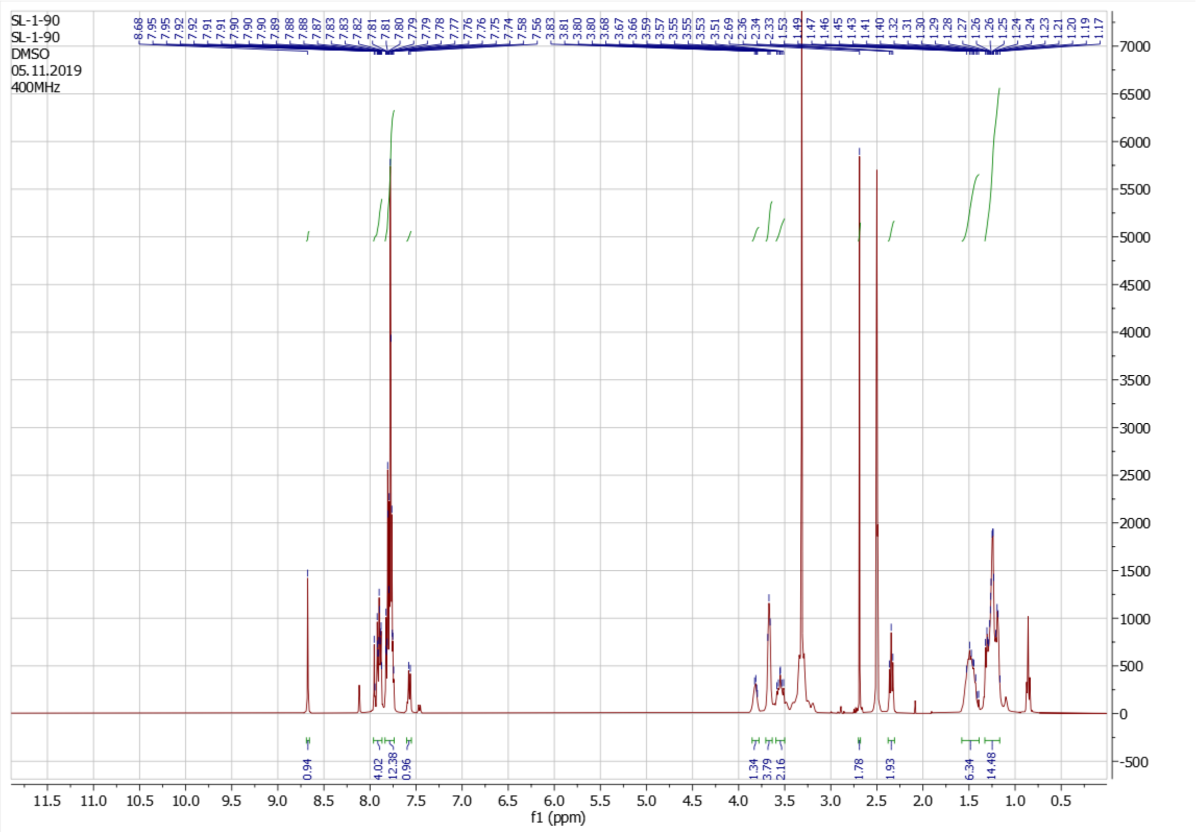

**Figure S15:** $^{1}$H NMR of **6** ON 400 MHz in DMSO-$d_6$.

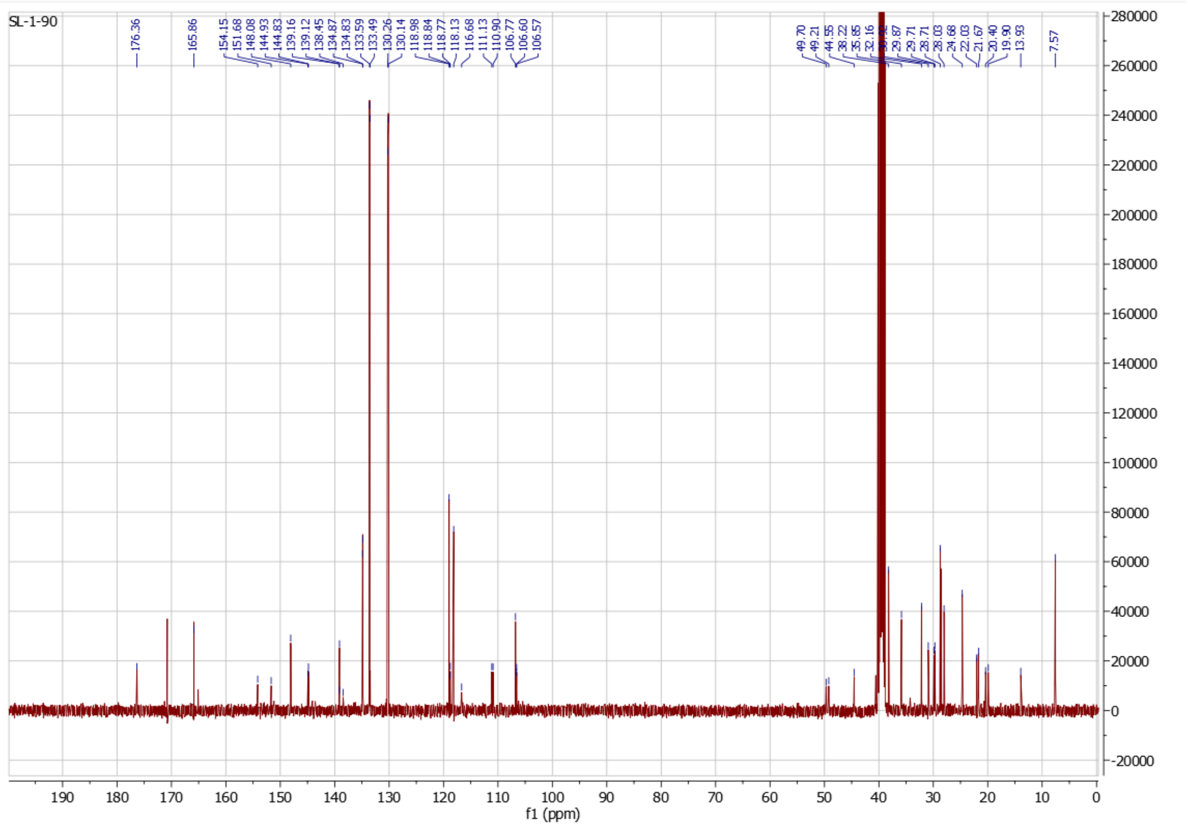

**Figure S16:** $^{13}$C NMR of **6** ON 101 MHz in DMSO-$d_6$.

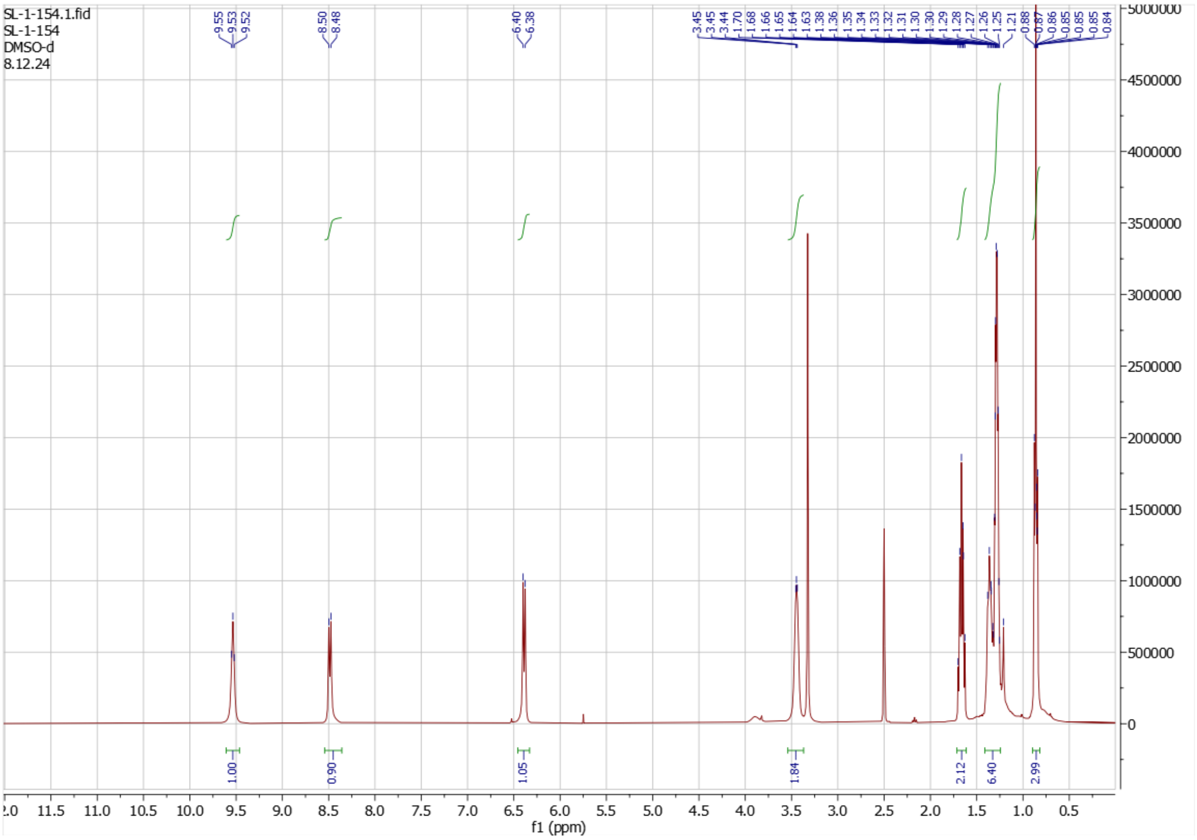

**Figure S17:** $^{1}$H NMR of **7** ON 400 MHz in DMSO-$d_6$.

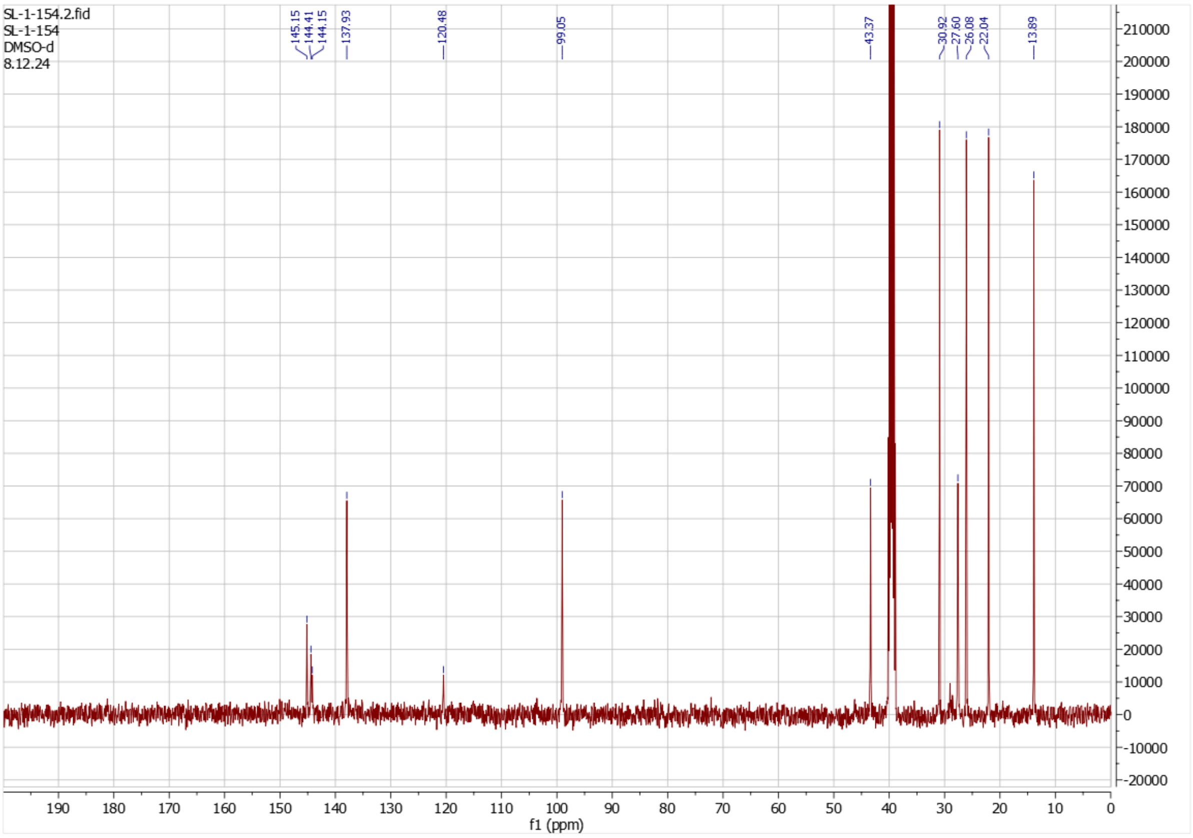

**Figure S18:** $^{13}$C NMR of **7** ON 101 MHz in DMSO-$d_6$.

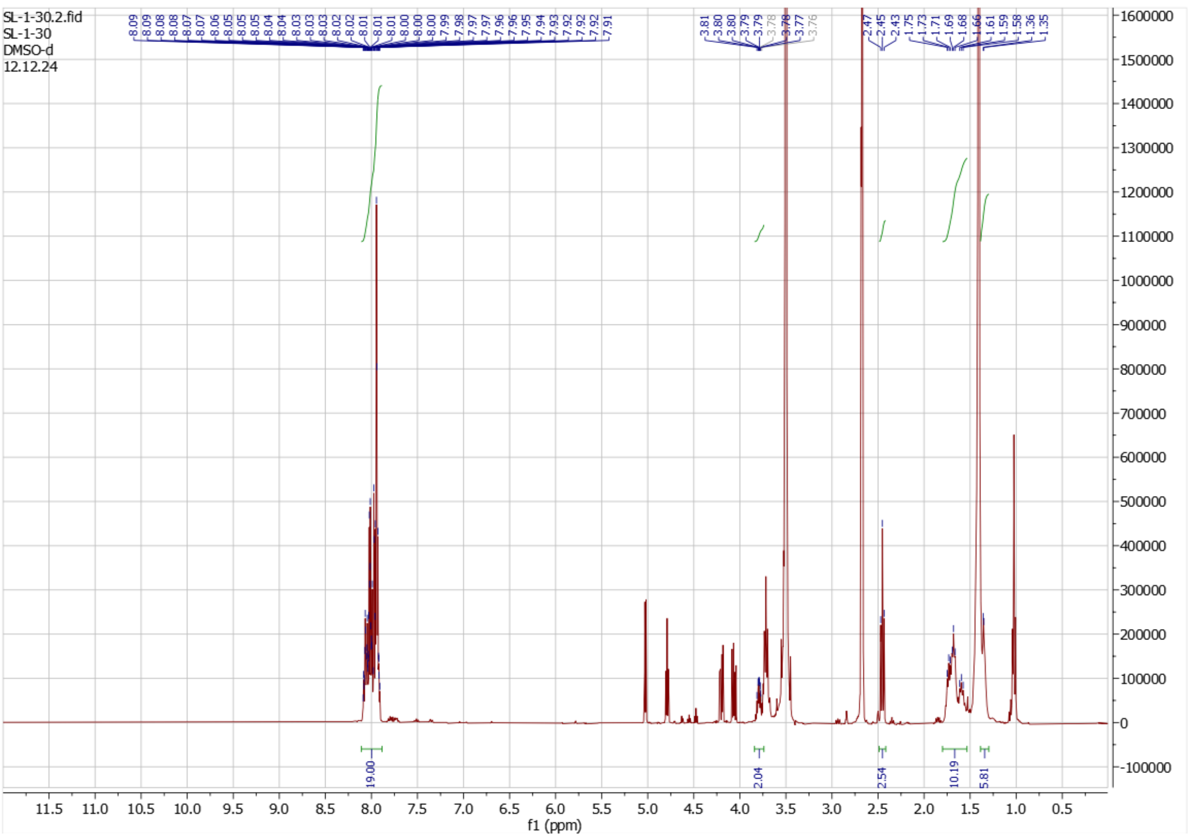

**Figure S19:** $^{1}$H NMR of **8** ON 400 MHz in DMSO-$d_6$.

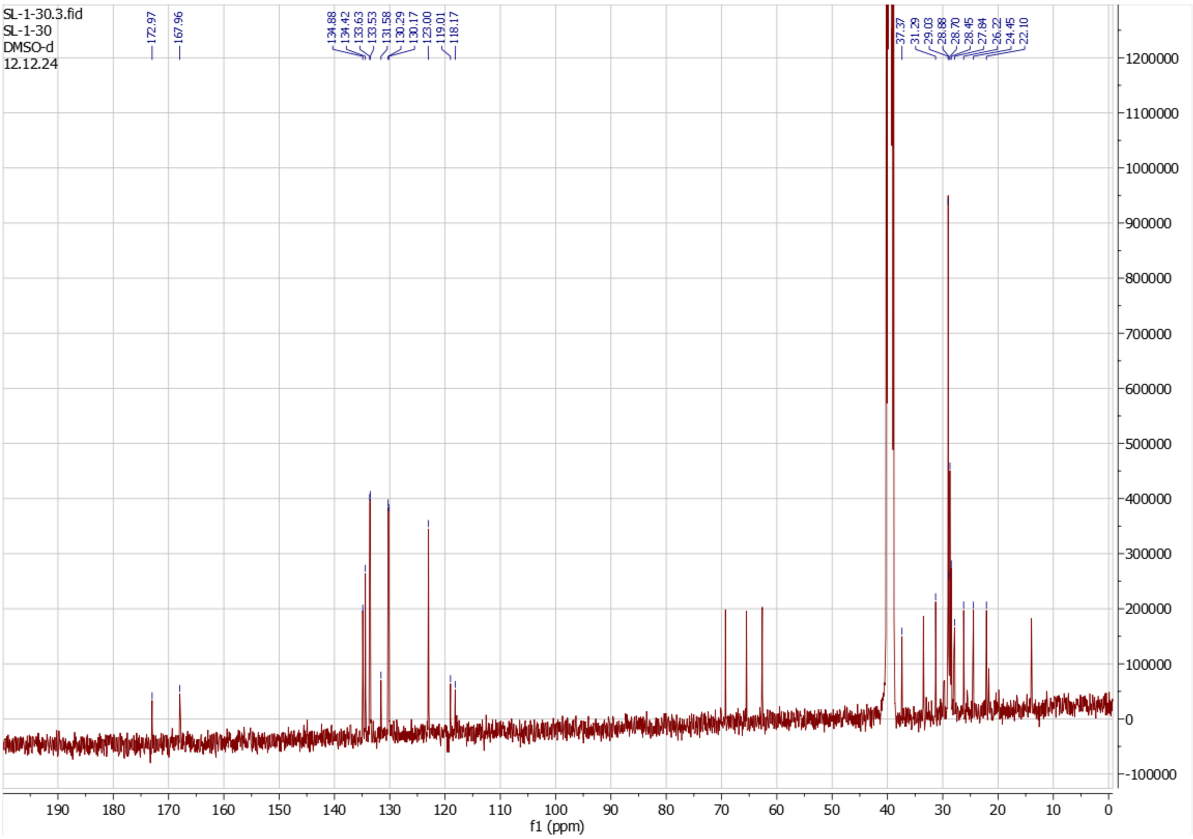

**Figure S20:** $^{13}$C NMR of **8** ON 101 MHz in DMSO-$d_6$.

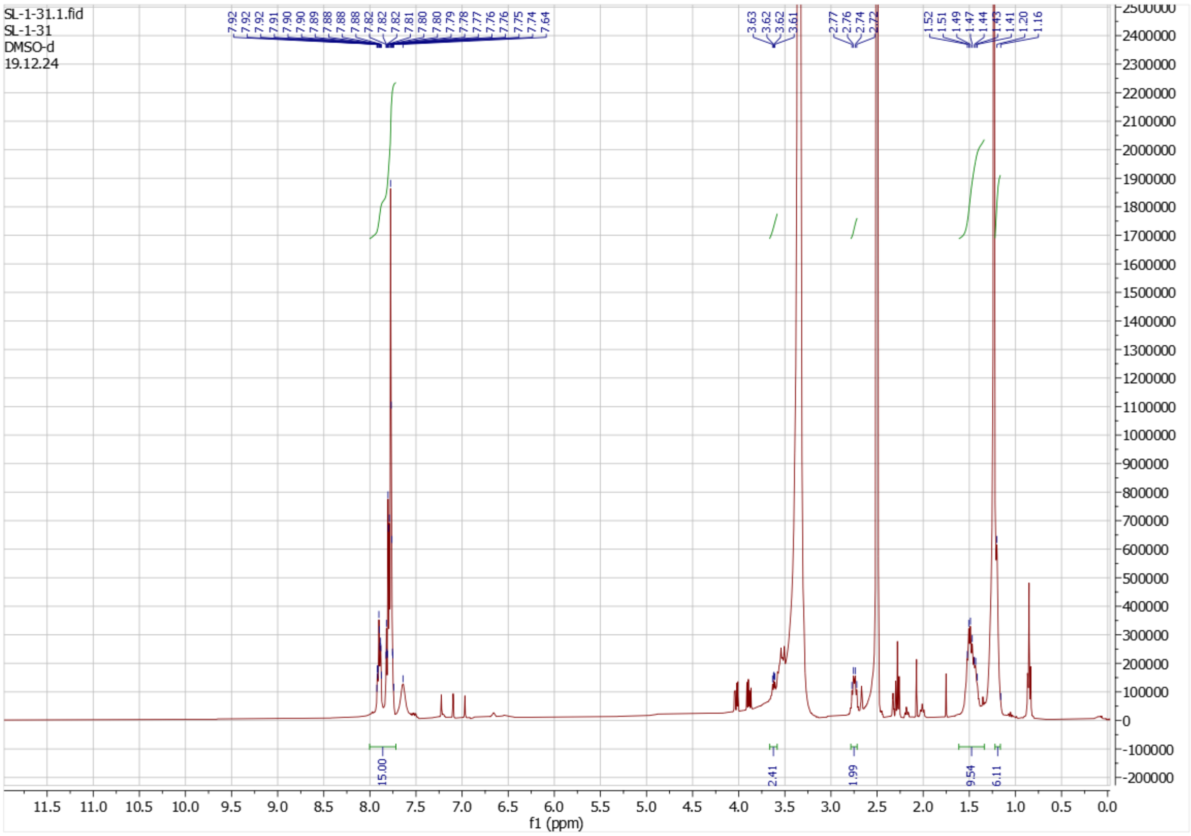

**Figure S21:** $^{1}$H NMR of **9** ON 400 MHz in DMSO-$d_6$.

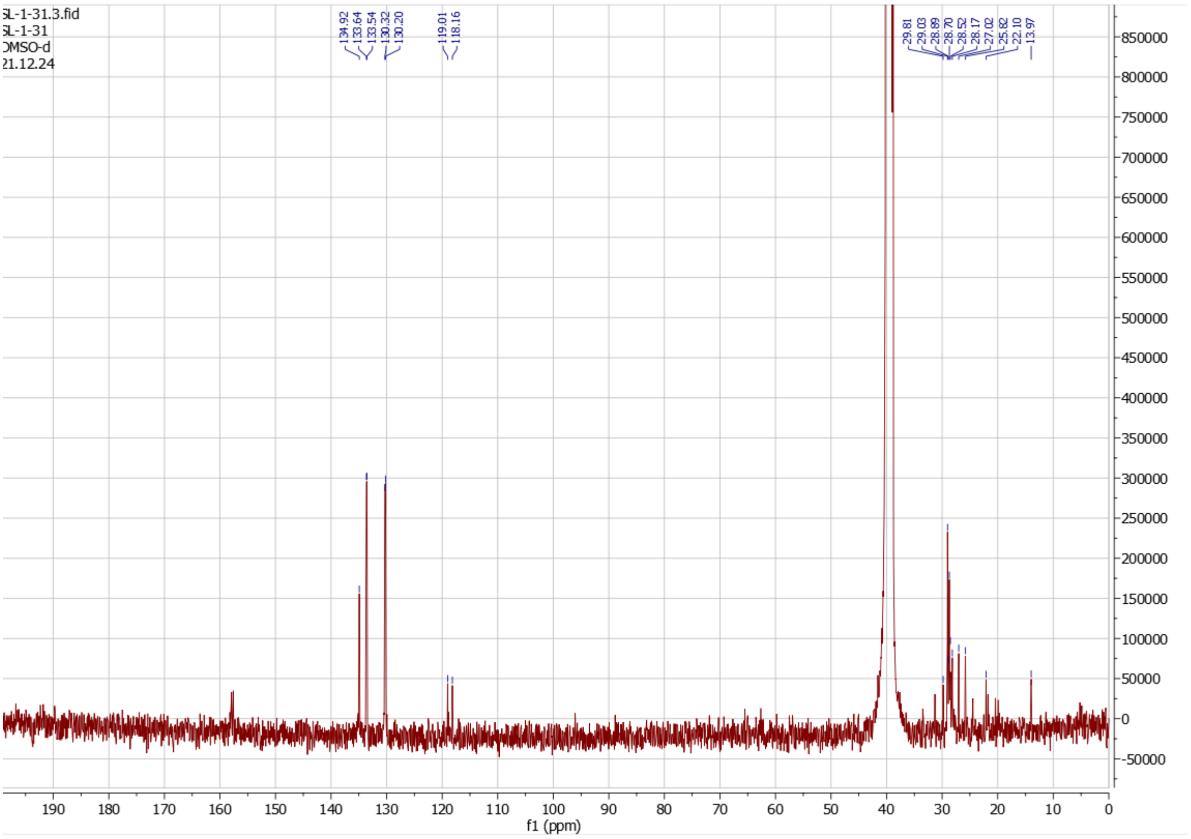

**Figure S22:** $^{13}$C NMR of **9** ON 101 MHz in DMSO-$d_6$.

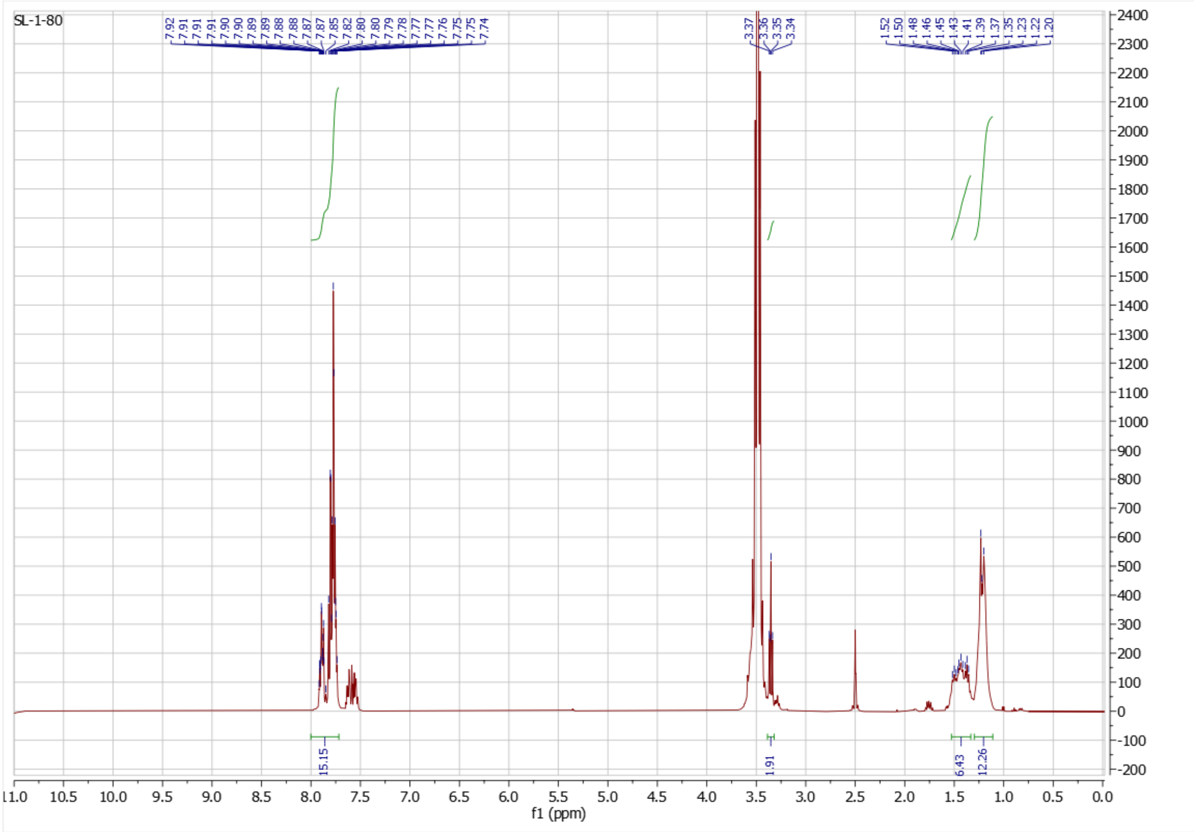

**Figure S23:** $^{1}$H NMR of **10** ON 400 MHz in DMSO-$d_6$.

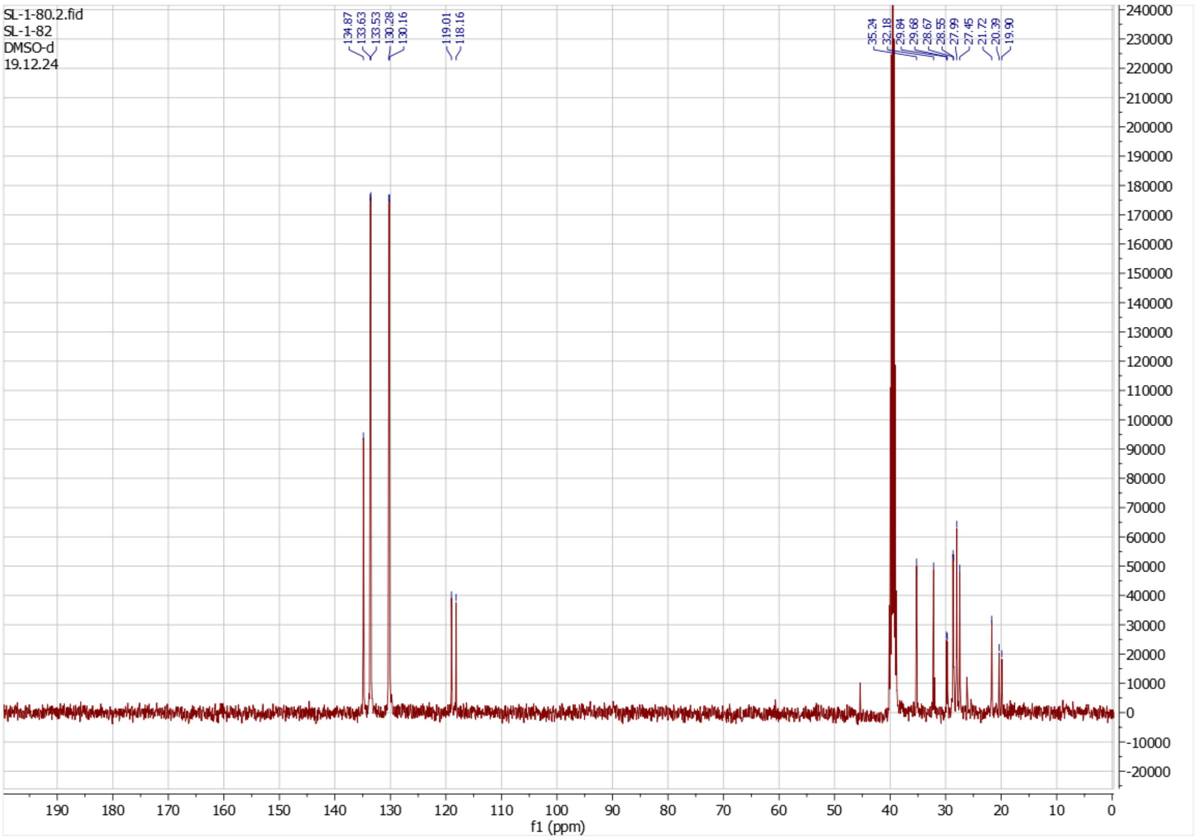

**Figure S24:** $^{13}$C NMR of **10** ON 101 MHz in DMSO-$d_6$.

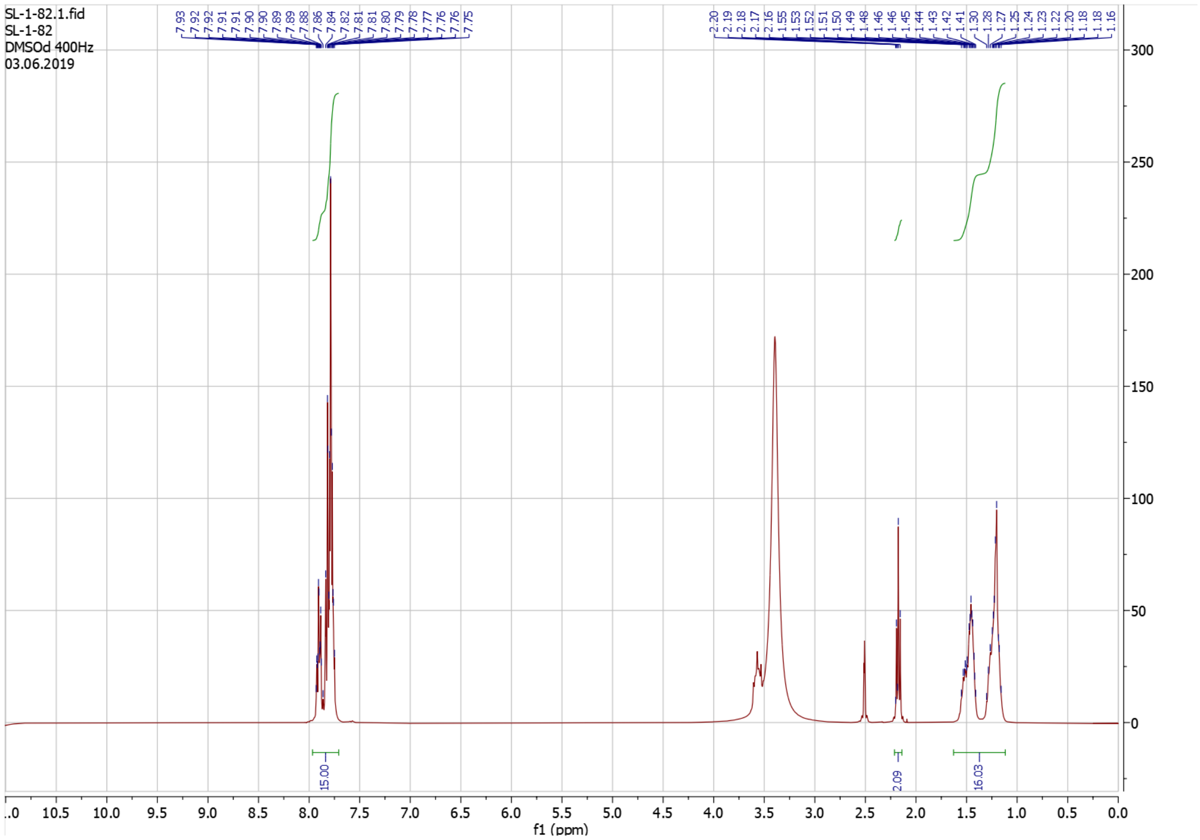

**Figure S25:** $^{1}$H NMR of **11** ON 400 MHz in DMSO-d$_6$.

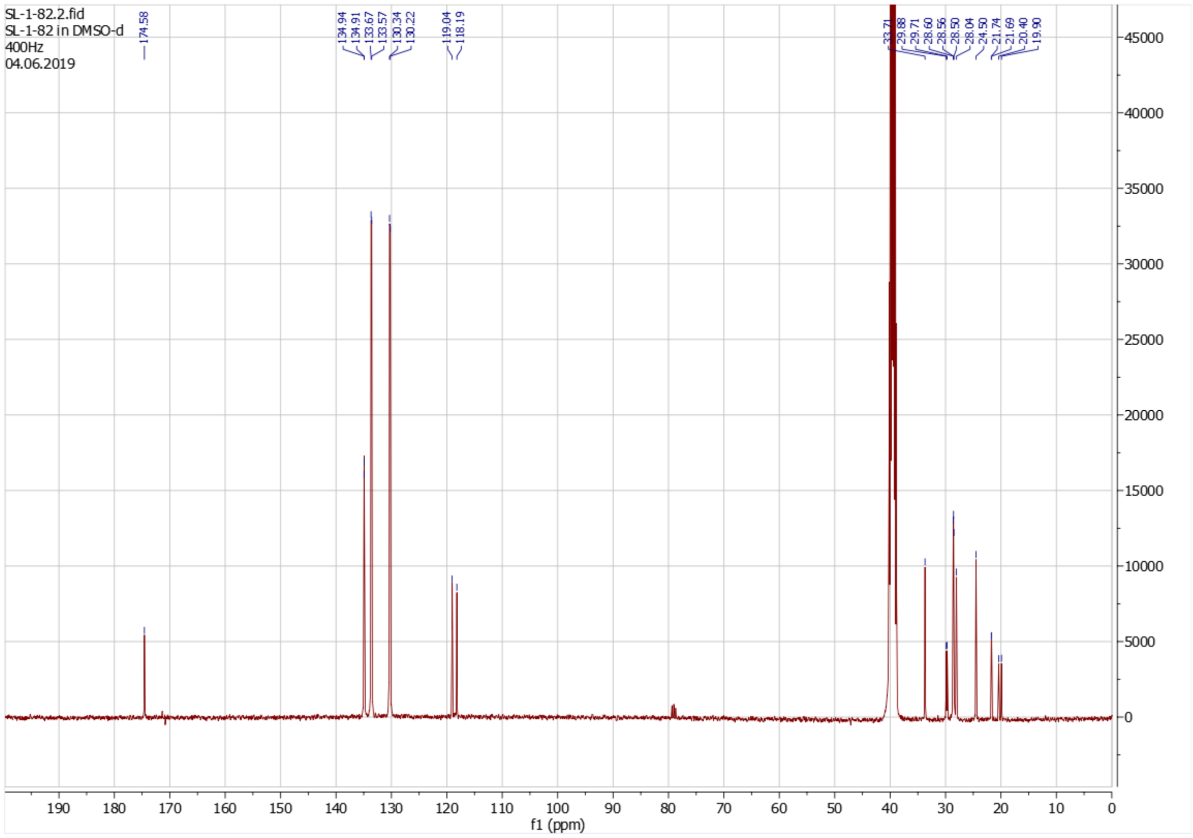

**Figure S26:** $^{13}$C NMR of **11** ON 101 MHz in DMSO-d$_6$.
